# Large scale functional screen identifies genetic variants with splicing effects in modern and archaic humans

**DOI:** 10.1101/2022.11.20.515225

**Authors:** Stephen Rong, Christopher R. Neil, Samantha Maguire, Ijeoma C. Meremikwu, Malcolm Meyerson, Ben J. Evans, William G. Fairbrother

## Abstract

Humans co-existed and interbred with other hominins which later became extinct. These archaic hominins are known to us only through fossil records and for two cases, genome sequences. Here we engineer Neanderthal and Denisovan sequences into thousands of artificial genes to reconstruct the pre-mRNA processing patterns of these extinct populations. Of the 5,224 alleles tested in this massively parallel splicing reporter assay (MaPSy), we report 969 exonic splicing mutations (ESMs) that correspond to differences in exon recognition between extant and extinct hominins. Using MaPSy splicing variants, predicted splicing variants, and splicing quantitative trait loci, we show that splice-disrupting variants experienced greater purifying selection in anatomically modern humans than in Neanderthals. Adaptively introgressed variants were enriched for moderate effect splicing variants, consistent with positive selection for alternative spliced alleles following introgression. As particularly compelling examples, we characterized a novel tissue-specific alternative splicing variant at the adaptively introgressed innate immunity gene *TLR1*, as well as a novel Neanderthal introgressed alternative splicing variant in the gene *HSPG2* that encodes perlecan. We further identified potentially pathogenic splicing variants found only in Neanderthals and Denisovans in genes related to sperm maturation and immunity. Finally, we found splicing variants that may contribute to variation among modern humans in total bilirubin, balding, hemoglobin levels, and lung capacity. Our findings provide novel insights into natural selection acting on splicing in human evolution and demonstrate how functional assays can be used to identify candidate causal variants underlying differences in gene regulation and phenotype.

## Introduction

Modern humans are the only present-day hominins, but used to co-exist with many others, including recently with our closest extinct archaic relatives, the Neanderthals and Denisovans. High-coverage ancient genomes sequenced from Neanderthal and Denisovan fossils have enabled the comparative study of genetic differences between modern and archaic humans (1–3). These genetic differences likely underlie many of the phenotypic differences between modern and archaic fossils, but also phenotypic differences not preserved in the fossil record (4, 5). Remarkably, genomic analyses have revealed multiple admixture events between modern and archaic humans (6–12), notably during modern human expansion Out of Africa. These introgression events contributed about 2% Neanderthal ancestry to all non-African populations (13), and up to 5% Denisovan ancestry to populations from Melanesia (11, 12, 14, 15). Because archaic humans had smaller effective population sizes than modern humans, they may have accumulated more deleterious alleles that, once introgressed, were subject to purifying selection in modern populations (16–18). Recent methods to detect introgressed segments (7–11, 19, 20) and adaptively introgressed loci (21–23) demonstrate a lasting impact of archaic introgression on several human phenotypes (4, 24), including those involved in local adaptation to novel environments (7, 9, 25), diets (26), and pathogens (27–30). Enrichment analyses of proteincoding and cis-regulatory regions show that introgressed variation has functional consequences in different tissue and cell types (1, 18, 31–33). One common approach is to look for trait-associated introgressed variants in modern human populations, such as those associated with complex traits (4, 24, 34–37), gene expression across tissues (38), or in response to infection (39–41). However, this approach is limited to studying introgressed variants found in well-sampled populations, which are currently biased towards individuals of European ancestry (42), and are ill-suited for studying modern or archaic-specific variants. Moreover, methods based on association tests are affected by allele frequency (AF) and linkage disequilibrium (LD) and often cannot distinguish causal variants from linked variants.

Recent studies have advanced several ways around these limitations. These include prediction models to impute unobserved differences in gene expression and 3D genome organization (43, 44); CRISPR-Cas9 editing of archaic alleles into human-derived stem cells (45, 46); and massively parallel reporter assays (MPRAs) to identify causal variants with cis-regulatory impacts on transcription (47–49). In particular, MPRAs have been applied to study modern-specific and adaptively introgressed variants (50–53). Most of these studies focused on transcriptional regulation even though post-transcriptional processes, such as RNA splicing regulation, are also important to human phenotypes and disease (54–57). Notable exceptions include an adaptively introgressed splicing QTL (sQTL) for isoforms of the *OAS1* gene involved in immune response (22, 28, 58, 59), a schizophrenia-associated introgressed variant that creates a novel splice acceptor site in the *ADAMTSL3* gene (38), and a survey of Neanderthal introgressed sQTLs involved in immune response in individuals of European ancestry (41).

Here, we analyzed a total of 5,224 modern, archaic, and introgressed variants in hominin evolution with a previously developed massively parallel splicing assay (MaPSy) (60). MaPSy is a parallel minigene reporter experimental system for identifying exonic splicing mutations (ESMs) that impact cis-elements involved in exon recognition during the process of pre-mRNA splicing. MaPSy and similar assays have been used to characterize splicing regulatory impacts of many types of variants, including pathogenic and rare variants (61–65). Like MPRAs for transcriptional regulation, MaPSy can distinguish causal from linked variants, and is not limited to variants observed in extant species. It is thus well-suited for characterizing the splicing effects of genetic variation within extant and extinct hominins. We use MaPSy-identified ESMs in conjunction with predicted splicing mutations and enrichment of sQTLs to characterize the relative contribution of splicing effects in hominin evolution. We considered possible effects of natural selection on these variants, and provide examples of candidate adaptively introgressed, lineage-specific, and introgressed trait-associated ESMs.

## Results

### Massively parallel splicing assay for identifying exonic splicing mutations

MaPSy identifies exonic splicing mutations (ESMs) with cis-regulatory effects on RNA splicing regulation in a high throughput minigene reporter experiment (Fig. 1a) (60). Briefly, a pool of oligos is synthesized containing pairs of ancestral (ANC) and derived (DER) endogenous DNA sequences for each single nucleotide variant (SNV) of interest. Every oligo contains a full exonic sequence with either the reference or alternative allele plus ≥55 bp of upstream and 15 of downstream intronic sequence. The oligo pool is then incorporated into a three-exon minigene reporter construct between common first and last exons with an upstream promoter and a downstream poly(A) site (Fig. S1a). The resulting pool of minigenes is then transfected in triplicate into human embryonic kidney (HEK) 293T cells, where they then undergo transcription followed by RNA splicing (Fig. S1b). Spliced RNA containing all three exons are mapped back to an allelic sequence. Deep resequencing of three input libraries of minigene DNA prior to transfection and three output libraries of cDNA from the spliced RNA was used to generate read counts for each allelic sequence. Read counts were highly replicable for both input DNA (*r* > 0.982) and output RNA (*r* > 0.991) libraries (Fig. S1c, d). Finally, mpralm (66) was used to estimate the allelic imbalance as the log_2_ fold change (FC) between DER/ANC reads in the output RNA/input DNA, along with p-values for the null hypothesis that log_2_ FC = 0. This allelic imbalance is the MaPSy functional score for the cis-effect on RNA splicing of the SNV. We defined ESMs as those variants with |log_2_FC| > log_2_(1.5) and an FDR-adjusted p-value < 0.05. We categorized ESMs as either promoting exon inclusion (positive ESMs) or inhibiting exon inclusion (negative ESMs), and as either strong ESMs (|log_2_FC| > log_2_(3)) or weak ESMs (log_2_(3) > |log_2_FC| > log_2_(1.5)). Typically, MaPSy experiments can only score ESMs in short exons (no more than 120 bp) due to length limitations of oligo synthesis (60). To account for additional variants in long exons, we validated a “half exon” approach to reliably score ESMs near the 3’ splice site (3’ ss) of longer exons (between 120-500 bp). See additional text in SI Appendix and Fig. S2.

**Figure 1.**
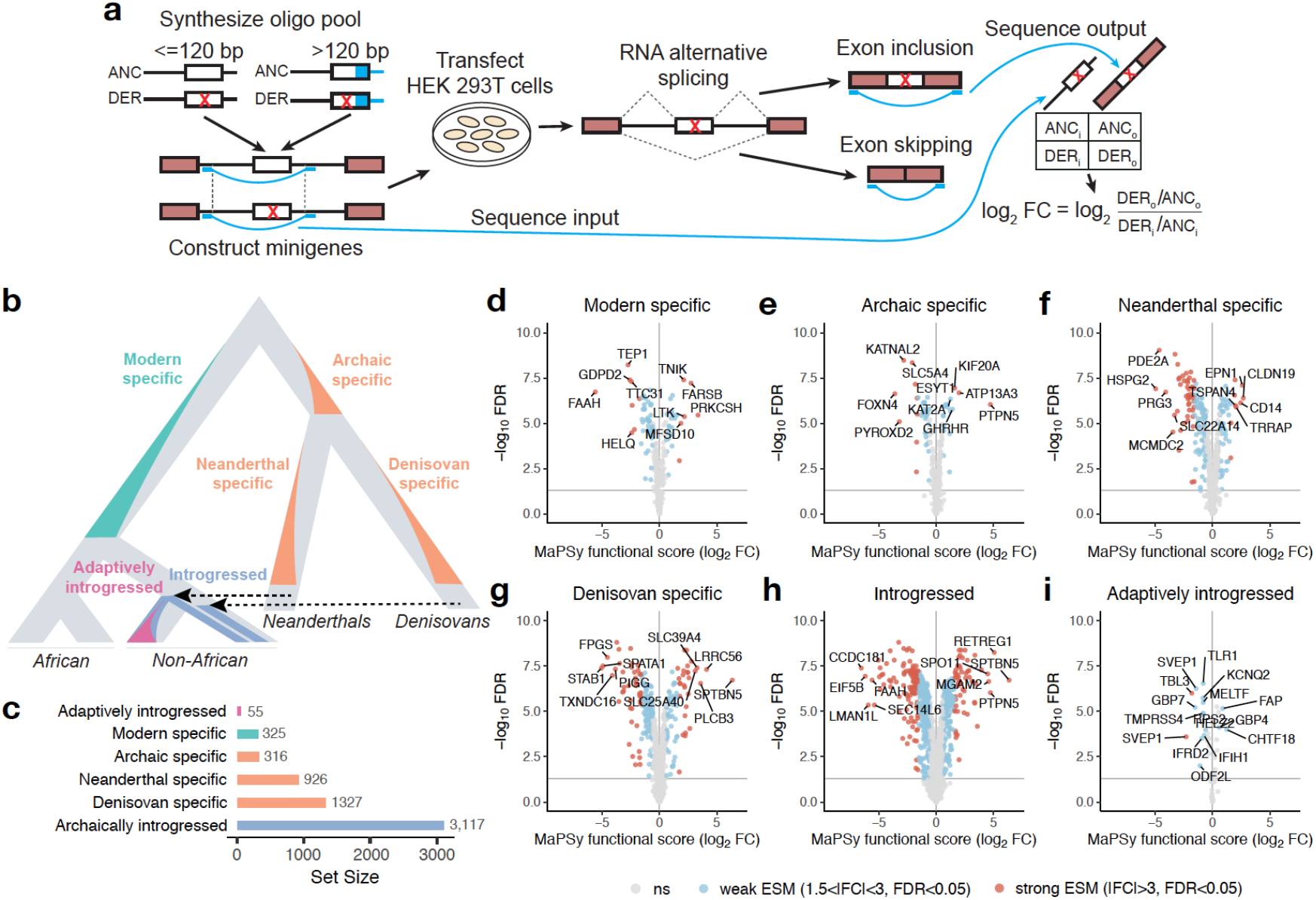
Massively parallel splicing assay for variants in human evolution. **(a)** Overview of the MaPSy experiment for identifying ESMs. **(b)** A simplified evolutionary tree showing the human evolution variant sets used in this study. **(c)** The number of variants from each variant set that were assayed in the MaPSy experiment. **(d-i)** MaPSy results for each of the human evolution variant sets. Each volcano plot shows the MaPSy functional score (log2 FC) versus the -log10 FDR (Benjamini-Hochberg-adjusted p-values, y-axis). Non-ESMS shown in grey, weak ESMs (1.5 < |FC| < 3) in blue, and strong ESMs (|FC| > 3) in red. Gene names for the top 10 ESMs ranked by |log2 FC| shown for each variant set, or for all ESMs for the adaptively introgressed variant set.

### Defining human evolution variant sets used in the MaPSy experiment

We defined six sets of human evolution-related variants based on comparisons between the 1000 Genomes Project (1KGP) (67), the Genome Aggregation Database (gnomAD) (68), four high-coverage archaic genomes, Altai Neanderthal, Vindija Neanderthal, Chagyrskaya Neanderthal, and Altai Denisovan (2, 15, 34, 69), and inferred human ancestral alleles from Ensembl (70) (Fig. 1b). The first four sets correspond to fixed or nearly fixed variants (i.e., above a threshold frequency as detailed below) with origins on different branches of the human evolutionary tree, which we refer to as X-specific variants for branch X. Modern-specific variants originated on the evolutionary branch leading to all modern humans, defined as having high frequency in modern humans (1KGP and gnomAD global AF > 0.9) and being absent in archaic humans (0/8 archaic alleles) in order to avoid incomplete lineage sorting variants. Archaic-specific variants originated on the branch between the split between modern and archaic humans but preceding the split between Neanderthals and Denisovans, defined as having high frequency in archaic humans (≥ 7/8 archaic alleles) and being nearly absent in modern African (AFR) populations (1KGP and gnomAD AFR AF < 0.01). Neanderthal-specific variants originated on the branch leading to Neanderthals after the split from Denisovans (≥ 5/6 Neanderthal alleles, 0/2 Denisovan alleles, 1KGP and gnomAD AFR AF < 0.01). Denisovan-specific variants originated on the branch leading to Denisovans after the split from Neanderthals (2/2 Denisovan alleles, 0/6 Neanderthal alleles, 1KGP and gnomAD AFR AF < 0.01). (See Methods and Fig. S3.) As only one genome was available, Denisovan-specific changes were not used outside the task of identifying example Denisovan-specific ESMs because there was less power to identify fixed or nearly fixed differences. Next, we defined an introgressed variant set, which contains previously published variants on archaically introgressed haplotypes from either Vernot et al. (8) or Browning et al. (11) identified in 1KGP populations and Papa New Guinea. Lastly, we defined an adaptively introgressed variant set, which contains previously published variants on high-frequency adaptively introgressed haplotypes from Gittelman et al. (22) and those on introgressed haplotypes from Browning et al. (11) overlapping high-frequency adaptively introgressed loci from Racimo et al. (23) in matching 1KGP populations. Some variants were found in multiple sets; for example, some archaic derived variants introgressed into modern humans (Fig. S4a,b). Modern-specific variants may overlap introgressed variants where introgression reintroduced the ancestral allele (51). Overall, we identified 2,036,206 variants belonging one of the human evolution variant sets (Additional file 2: Table S1), of which 5,224 exonic variants could be incorporated into the MaPSy experiment (Fig. 1c and Additional file 3: Table S2).

### MaPSy identifies hundreds of ESMs in human evolution

We applied MaPSy to the 5,224 human evolution variants and identified 969 ESMs (19% of assayed variants) that were significant at an FDR threshold of < 0.05 and a MaPSy functional score threshold of |FC| > 1.5. This included 71 (of 325) modern-specific, 51 (of 316) specific to the most recent common ancestor of Neanderthals and Denisovans, hereafter archaic-specific, 170 (of 926) Neanderthal-specific, 232 (of 1,327) Denisovan-specific, 591 (of 3,117) introgressed, and 16 (of 55) adaptively introgressed ESMs (Fig. 1d–i). 683 ESMs (13%) have weak effects on splicing (weak ESMs), and 286 ESMs (5%) have strong effects (strong ESMs).

### Weak and strong ESMs affect splicing through distinct mechanisms

Many ESMs were found in splice region positions closest to the 3’ ss and 5’ ss, particularly ESMs with the strongest negative effects on splicing (strong negative ESMs) that were expected to result in loss of canonical transcript exon inclusion (71) (Fig. S5a). The vast majority of ESMs, however, were synonymous and missense mutations found deeper into the exon with relatively moderate positive or negative effects on exon inclusion (weak ESMs). To better understand the mechanisms by which ESMs affect splicing, we compared MaPSy functional scores to computational predictions of different splicing effects. First, we used SpliceAI, a deep neural network model trained to predict the location of splice sites from RNA sequence, which can be used to identify mutations that disrupt splice sites (72). We used the max of raw SpliceAI scores for predicted donor gain, donor loss, acceptor again, and acceptor loss (SpliceAI max score). SpliceAI max scores tend to increase with MaPSy scores away from zero, and the variants with the highest SpliceAI scores tend to also have the most negative (but not positive) MaPSy scores (Fig. 2a). Next, we used Delta EI scores, which is the change in the ability of overlapping hexamers at a SNV to function as exonic splicing enhancers (73). SNVs with positive/negative Delta EI are predicted to increase/decrease exon inclusion. Consistent with this, we find that Delta EI scores increase with MaPSy scores, except when MaPSy scores are extremely negative (Fig. 2a). We find similar trends for the change in two alternative hexamer scores of exonic splicing enhancer activity (74) (Fig. S5b,c). Overall, these results suggest that MaPSy is sensitive at detecting ESMs acting through distinct splicing mechanisms. First, those with strong negative effects (strong negative ESMs) on splicing tend to be found in splice regions, affect splicing through the mechanism of splice site disruption, and are thus likely to be deleterious. Second, those with weaker effects on splicing (weak ESMs) are enriched in the interior of the exon, and affect splicing through changes in exonic splicing regulatory motifs.

**Figure 2.**
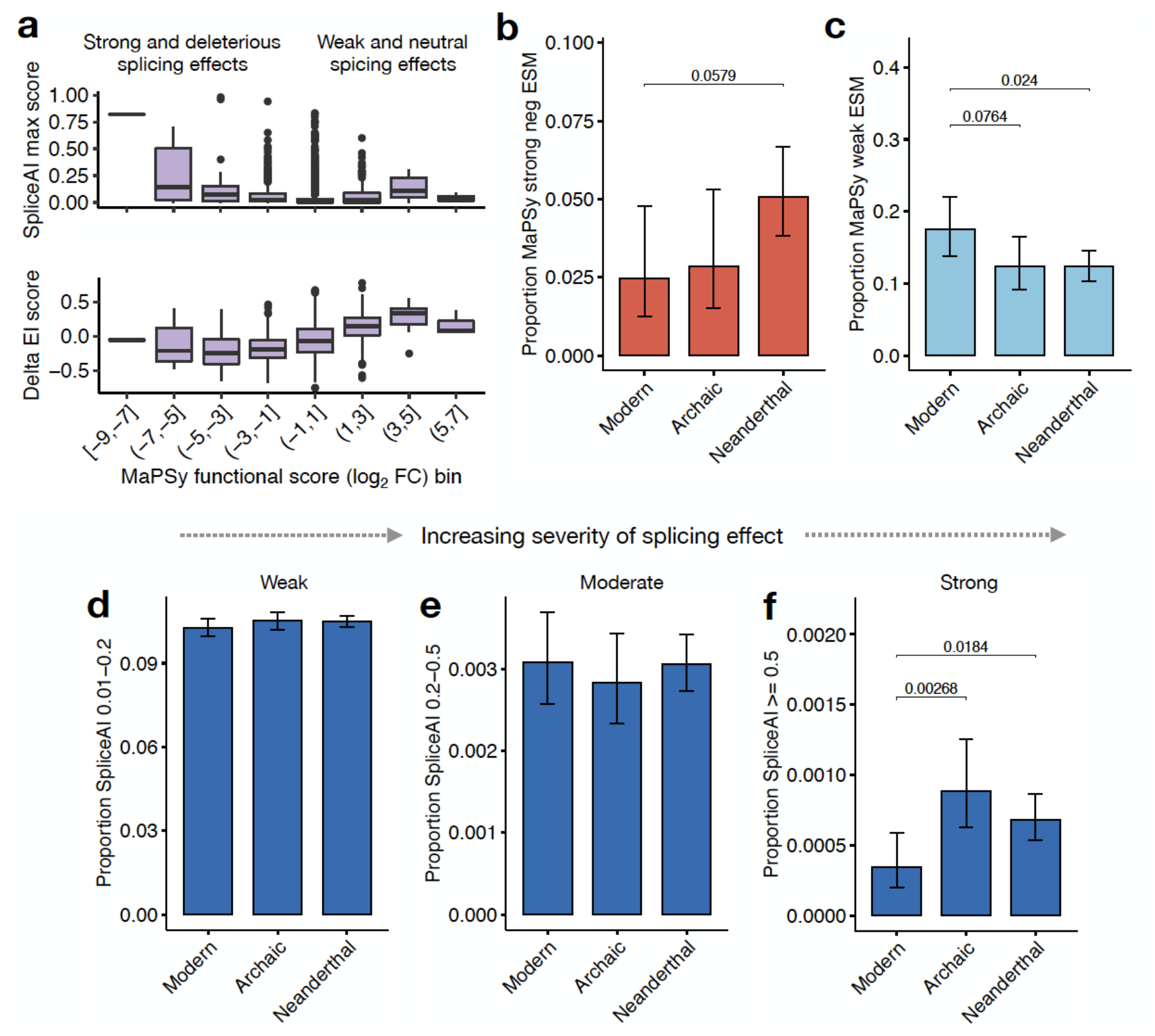
Modern human lineage is depleted in strong splicing effects relative to the archaic human lineage. **(a)** Relationship between MaPSy functional score (log2 FC) bins versus SpliceAI max score and Delta EI scores. **(b-c)** Differences in strong negative (FC < −3) and weak (1.5 < |FC| < 3) ESM proportions between modern-, archaic-, and Neanderthal-specific variants. **(d-f)** Variants were binned by SpliceAI max raw scores into no splicing effect (not shown), weak splicing effect (between 0.01 and 0.2), moderate splicing effect (0.2-0.5), and strong splicing effect (0.5-1). Variant sets were compared based on the proportion of variants in SpliceAI bins between modern-, archaic-, and Neanderthal-specific variants. Pairwise P-values from Fisher’s exact tests, p-values < 0.1 shown.

### Modern humans are depleted in strong splicing effects relative to archaic humans

Purifying selection is expected to be more efficient in modern humans as compared to Neanderthals and Denisovans owing to the larger effective population size of modern humans. We therefore reasoned that purifying selection might affect the distribution and function of ESMs in different hominin populations (16–18). We tested this hypothesis by comparing proportions of fixed or nearly fixed strong negative MaPSy ESMs and strong SpliceAI scores specific to the modern, archaic, and Neanderthal branches. We focused on these variants because strong negative, but not positive, MaPSy ESMs and strong SpliceAI scores are in general depleted in evolutionarily constrained genes (Fig. S6) as defined by low loss-of-function observed/expected upper bound fraction (LOEUF) in gnomAD (75). Modern-specific variants were found to be nominally depleted in strong negative ESMs compared to Neanderthal-specific variants (*p* = 0.0579) (Fig. 2b), but not in strong positive ESMs (Fig. S7a). This is consistent with stronger purifying selection in modern versus archaic humans. They were also enriched in weak ESMs (*p* = 0.024) (Fig. 2c). Considering a range of SpliceAI scores of increasing severity of splicing effect from weak (0.01-0.2), to moderate (0.2-0.5), to strong (0.5-1.0), modern-specific variants were significantly depleted in strong SpliceAI scores compared to both archaic-specific (*p* = 0.00268) and Neanderthal-specific variants (*p* = 0.0184) (Fig. 2f), concordant with the MaPSy results. However, all three branches had similar proportions of weak and moderate SpliceAI scores (Fig. 2d,e). Together, the MaPSy and SpliceAI results support the hypothesis that modern humans are depleted in ESMs strongly associated with deleterious exon skipping as compared to Neanderthals. This is consistent with expectations based on their relative effective population sizes. Concordant with this, we also observed a similar depletion of missense variants in the modern lineage compared to the archaic (*p* = 0.00868) and Neanderthal (*p* = 0.00403) lineages, but not of synonymous (as expected) nor nonsense and splice region variants (likely due to low statistical power) (Fig. S9). No differences in proportions of weak, moderate, or strong SpliceAI variants were found across modern human populations in the 1KGP (based on variants with MAF ≥ 0.01, Fig. 8), which is consistent with 1KGP modern Africans and non-Africans having much higher heterozygosity than archaic humans (2, 15, 34).

### Adaptively introgressed variants are enriched in moderate splicing effects relative to all introgressed variants

Introgressed variants in modern human populations are known to be depleted in genes, conserved coding regions, and non-coding regulatory regions, which could be due to purifying selection against deleterious introgressed alleles (9, 18, 31–33). However, high-frequency adaptively introgressed variants are known to be enriched in genes and non-coding regulatory regions, which is likely due to positive selection (31, 33). Supporting this, many adaptively introgressed haplotypes have been shown to affect gene expression (22, 28, 29, 39, 40). We tested the hypothesis that adaptively introgressed variants are enriched in splicing effects compared to all introgressed variants using both MaPSy scores and SpliceAI scores. Adaptively introgressed variants were found to be enriched in weak ESMs compared to all introgressed variants (*p* = 0.0161) (Fig. 3a), but not in strong negative or strong positive ESMs (Fig. S7b,c). We considered a range of SpliceAI scores from weak (0.01-0.2), to moderate (0.2-0.5), to strong (0.5-1.0). Adaptively introgressed variants were found to be significantly enriched in moderate SpliceAI scores compared to all introgressed variants (*p* = 0.0433) (Fig. S10b), but not weak or strong SpliceAI scores (Fig. S10a,c), which is concordant with the MaPSy results. Together, these findings suggest that adaptively introgressed variants are enriched in splicing variants that may be involved in alternative splicing events compared to all introgressed variants.

**Figure 3.**
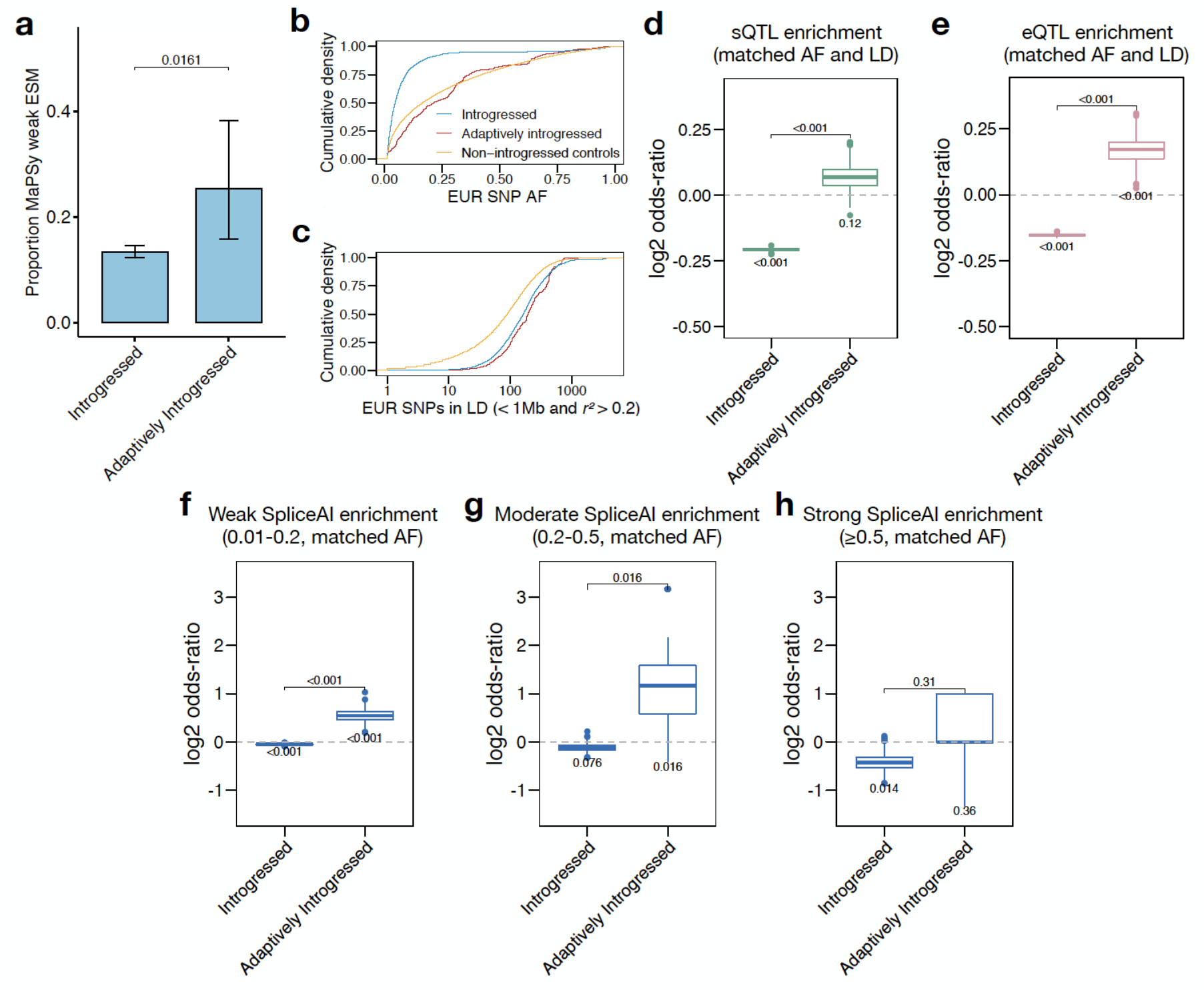
Introgressed variants underlying local adaptation are enriched in moderate splicing effects. **(a)** Differences in weak (1.5 < |FC| < 3) ESM proportions between introgressed and adaptively introgressed variants. Pairwise P-values from Fisher’s exact tests, p-values < 0.1 shown. **(b-c)** Allele frequency (AF) and linkage disequilibrium (LD) differences between introgressed, adaptively introgressed, and non-introgressed variants in 1KGP European populations, as represented by cumulative densities of 1KGP European AF (EUR SNP AF) and a univariate measure of LD (the number of EUR SNPs in LD within 1 Mb with *r^2^* > 0.2). **(d-e)** Enrichment of GTEx sQTLs and eQTLs significant at FDR < 0.05 in at least one tissue among introgressed and adaptively introgressed variants. Variability of estimates, one-sample p-values, and pairwise p-values are based on 1,000 randomly sampled sets of AF and LD matched controls. See methods for details on p-value calculations. **(f-g)** Enrichment (log2 odds-ratio) relative to AF matched controls among introgressed and adaptively introgressed variants for weak SpliceAI scores (max raw score between 0.01-0.2), moderate SpliceAI scores (0.2-0.5), and strong SpliceAI scores (0.5-1). Variability of estimates, one-sample p-values, and pairwise p-values are based on 1,000 randomly sampled sets of AF matched controls.

### Purifying selection on introgressed and positive selection on adaptively introgressed splicing variation

The pattern of ESM enrichment detailed above could stem from positive selection favoring beneficial splicing variants or from purifying selection disfavoring deleterious splicing variants. We thus sought to independently evaluate enrichment of splicing variants among the introgressed or adaptively introgressed variant sets relative to matched non-introgressed control variants. For splicing variants, we used Genotype Tissue-Expression (GTEx) sQTLs and variants with weak (0.01-0.2), moderate (0.2-0.5), and strong (0.5-1) SpliceAI scores. Since archaic introgression itself generates LD, introgressed variants are more likely to tag sQTLs, and thus it is necessary to use AF and LD matched non-introgressed controls to assess enrichment of sQTLs (Fig. S11a,b). Compared to non-introgressed variants, introgressed variants have overall lower AF and higher LD, whereas adaptively introgressed variants have similar AF but higher LD (Fig. 2b,c). We thus measured enrichment relative to randomly sampled, non-introgressed variants in matched AF and LD bins. This approach showed that we could recapitulate known intergenic enrichment and missense depletion among introgressed variants, and genic enrichment among adaptively introgressed variants (Fig. S11c,d).

In analyzing QTL enrichment, we found that introgressed variants were significantly depleted in GTEx-based sQTL variants (those with FDR < 0.05 in at least one tissue) (*p* < 0.001), adaptively introgressed variants are nominally enriched in QTLs (*p* = 0.12), and there is a significant difference between the two sets (*p* < 0.001) (Fig. 3d). We found a similar pattern for enrichment of overlap with GTEx-based expression QTLs (eQTLs), though introgressed variants were more depleted for sQTLs than eQTLs, and adaptively introgressed variants were more enriched for eQTLs than sQTLs (Fig. 3e). We also found introgressed variants were most depleted in strong SpliceAI scores (*p* = 0.014) (Fig. 3h). Adaptively introgressed variants were enriched in weak (*p* < 0.001) and moderate SpliceAI scores (*p* = 0.016), but not strong SpliceAI scores (*p* = 0.36) (Fig. 3f,g). Overall, these ESM, sQTL, and SpliceAI enrichment analyses suggest introgressed variants are depleted in variants that result in loss of exon inclusion, which is consistent with the action of purifying selection post-introgression. In contrast, adaptively introgressed variants are enriched in moderate effect splicing variants that change splicing relative to the hominin ancestor, suggesting the action of positive selection post-introgression. Possible explanations for why we did not observe an enrichment of strong effect splicing variants in adaptive introgression are provided below.

### MaPSy identifies a causal variant in the adaptively introgressed TLR1 sQTL

As discussed above, linkage disequilibrium (LD) often limits the ability to identify an individual variant’s contribution to a QTL, but MaPSy provides a functional test of the splicing phenotype of an individual nucleotide variant. To better understand the contribution of introgressed variants to an sQTL, MaPSy hits were compared to sQTL blocks and predictive measures of splicing effect such as Splice AI prediction. We showed above that introgressed variants overall were depleted in strong negative ESMs, strong SpliceAI scores, and sQTLs, suggesting a role of purifying selection against splicing disruptions. However, the subset of adaptively introgressed variants showed enrichment of moderate splicing effects. One such example is an adaptive Neanderthal-introgressed ESM, rs5743566, in the Toll-like receptor gene *TLR1* involved in innate immunity (Fig. 4). The SNV rs5743566 belongs to an adaptive Neanderthal-introgressed haplotype spanning the *TLR6/TLR1/TLR10* locus, a hotspot of repeated introgression from both Neanderthals and Denisovans (29). The introgressed haplotypes at this locus have previously shown to have eQTL effects on each of *TLR1, TLR6*, and *TLR10* (22, 29, 38) and sQTL effects on *TLR1* (41) in response to infection. The SNV rs5743566 resides in non-coding exon 2 of *TLR1* in the 5’ UTR region 15 bp away from the nearby 5’ ss and has a MaPSy functional score of −0.81, suggesting the derived G mutation moderately reduces inclusion of the exon compared to the ancestral C allele. It has a SpliceAI donor loss score of 0.38 affecting the nearby 5’ ss, which is the highest SpliceAI score of any variant on the adaptively introgressed haplotype (Fig. 4a). An analysis of GTEx sQTLs shows rs5743566 has strong association with *TLR1* splicing phenotype in 7 different tissues and resides in a large sQTL block containing dozens of other associated variants in LD (Fig. 4a). Comparisons of genotypes in three Neanderthal and one Denisovan genomes confirms that rs5743566 likely originated in and was nearly fixed in the Neanderthal population (Fig. 4b), and was later adaptively introgressed into modern human populations where it is now found at intermediate frequencies in non-African populations in 1KGP (Fig. 4c). Finally, a motif-based search for RNA binding protein (RBP) motifs using the ATtRACT database (76) suggests the C>G mRNA change introduces a UG-rich binding site for CELF proteins, a family of RBPs known to regulate alternative splicing (77) (Fig. 4d). This is consistent with a Delta EI score −0.22 for the C>G mRNA change suggesting a moderate loss of exonic splicing enhancer activity. This example evidences the ability of MaPSy to identify causal splicing variants in an adaptively introgressed sQTL block.

**Figure 4.**
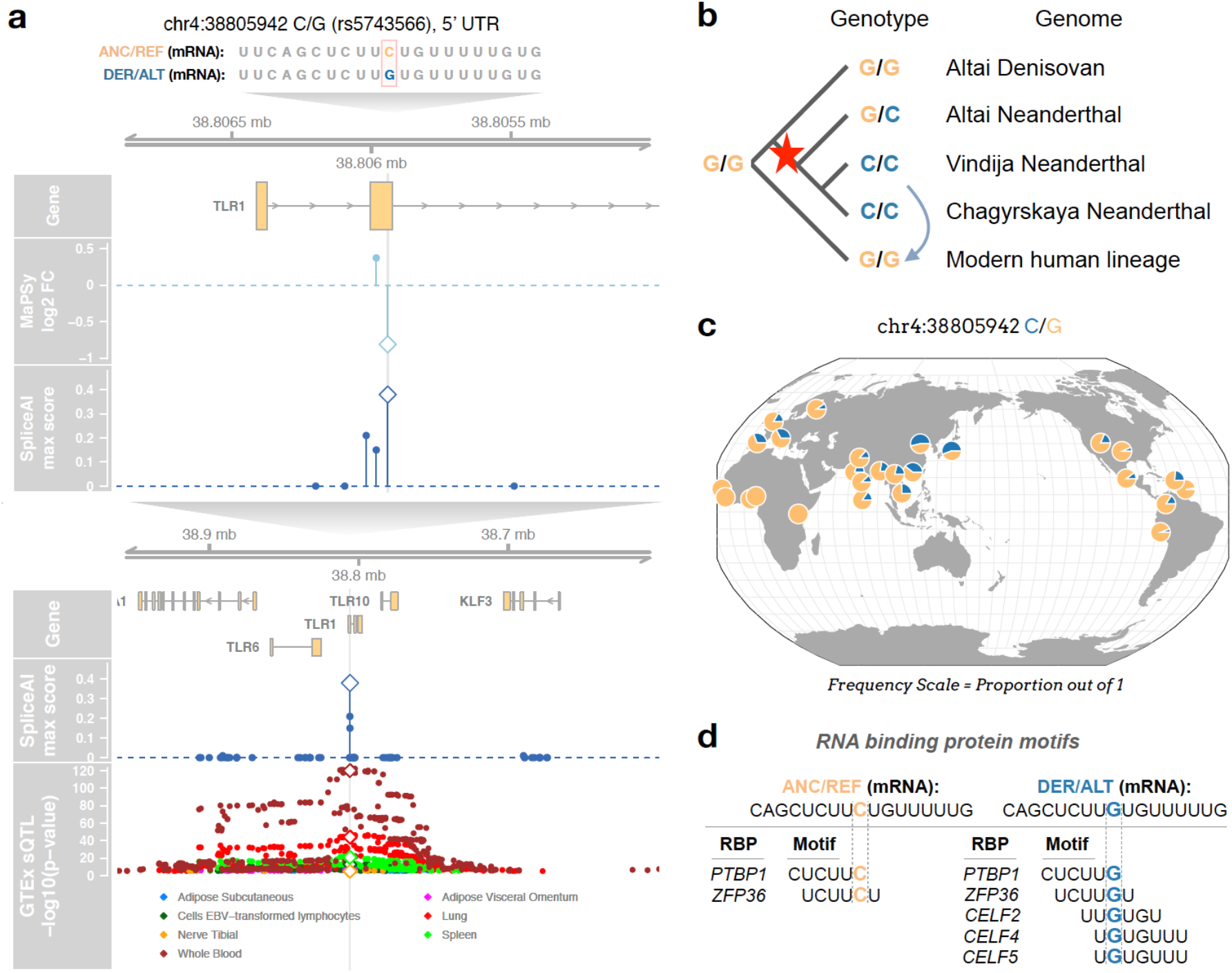
Adaptively introgressed ESM (rs5743566) in the innate-immunity gene *TLR1*. **(a)** Genome tracks showing MaPSy functional scores (log2 FC), SpliceAI max scores, and GTEx sQTL results for rs5743566 (diamonds) and nearby introgressed variants. GTEx sQTL scores are -log10 p-values for all SNVs that are significant *TLR1* sQTLs at FDR < 0.05 in at least one tissue. The variant rs5743566 likely arose in Neanderthals and was subsequently adaptively introgressed into modern human populations, as supported by **(b)** comparisons between genotypes in four high-coverage archaic genomes and the consensus modern human genotype (red star represents origin of variant, light blue arrow represents introgression event), and **(c)** 1KGP allele frequencies across 26 human populations from the Geography of Genetic Variants Browser. **(d)** RNA binding protein (RBP) motifs (min width of 6) identified in the ANC or DER sequences using ATtRACT.

### MaPSy identifies lineage-specific causal splicing variants

We next considered as case-studies the strongest Neanderthal-specific, archaic-specific, and modern-specific ESMs that were identified by their MaPSy functional scores (Additional file 4: Table S3). The strongest Neanderthal-specific ESM that was also introgressed was rs12737091, a synonymous variant in the gene *HSPG2. HSPG2* encodes the protein perlecan, a key component of the extracellular matrix that has diverse roles, including contributing to cartilage and bone development and angiogenesis (78). This variant has a MaPSy function score of −4.95 and a SpliceAI donor gain score of 0.98, which is consistent with the C>A mRNA change creating a GUAAG canonical splice donor motif, and is a sQTL for *HSPG2* in thyroid, adipose (subcutaneous), and artery (tibial) tissues (Fig. 5a). The derived allele is fixed in all three Neanderthal genomes, and is found at low frequencies in non-African modern humans (Fig. 5b,c). The strongest archaic-specific ESM that was also introgressed was rs17518147 in the splice region of exon 2 of *CCDC181*, a gene that encodes a microtubule-binding protein in sperm (Fig. S12) (79). It has a MaPSy functional score of −6.53 and a SpliceAI donor loss score of 0.59, suggesting the G>A mRNA change negatively disrupts the adjacent splice donor site (Fig. S12a). The derived allele is fixed across all four archaic genomes, suggesting an origin prior to the split between Neanderthals and Denisovans (Fig. S12b), and was later introgressed from Neanderthals into modern humans where it is now found at low to intermediate frequencies across non-African populations (Fig. S12c). It is also identified in GTEx as a *CCDC181* sQTL in testis tissue. The strongest modern-specific ESM was rs77724956 in the splice region of exon 12 of *FAAH*, a gene that encodes a catabolic enzyme for fatty acid amides such as anandamide involved in pain sensation (Fig. S13) (80). It has a MaPSy functional score of −5.59 and a SpliceAI acceptor loss score of 0.28, suggesting the G>U mRNA change results in a negative disruption of the adjacent splice acceptor (Fig. S13a). The derived T allele is not found in any of the four archaic genomes, and is nearly fixed in modern humans worldwide (Fig. S13b). The ancestral G allele is found at a very low frequency in modern humans, with the highest population frequency of 1-2% in East and South Asia (Fig. S13c). Notably, rs77724956 is also an introgressed variant, suggesting the functional ancestral allele was reintroduced into modern human populations (51). It is nearly absent in European populations, and thus has no associated GTEx sQTL results. However, the eQTLGen consortium, which provides whole blood eQTLs based on 31,684 individuals across multiple ancestries, identifies rs77724956 as a strongly associated *FAAH* eQTL (81).

**Figure 5.**
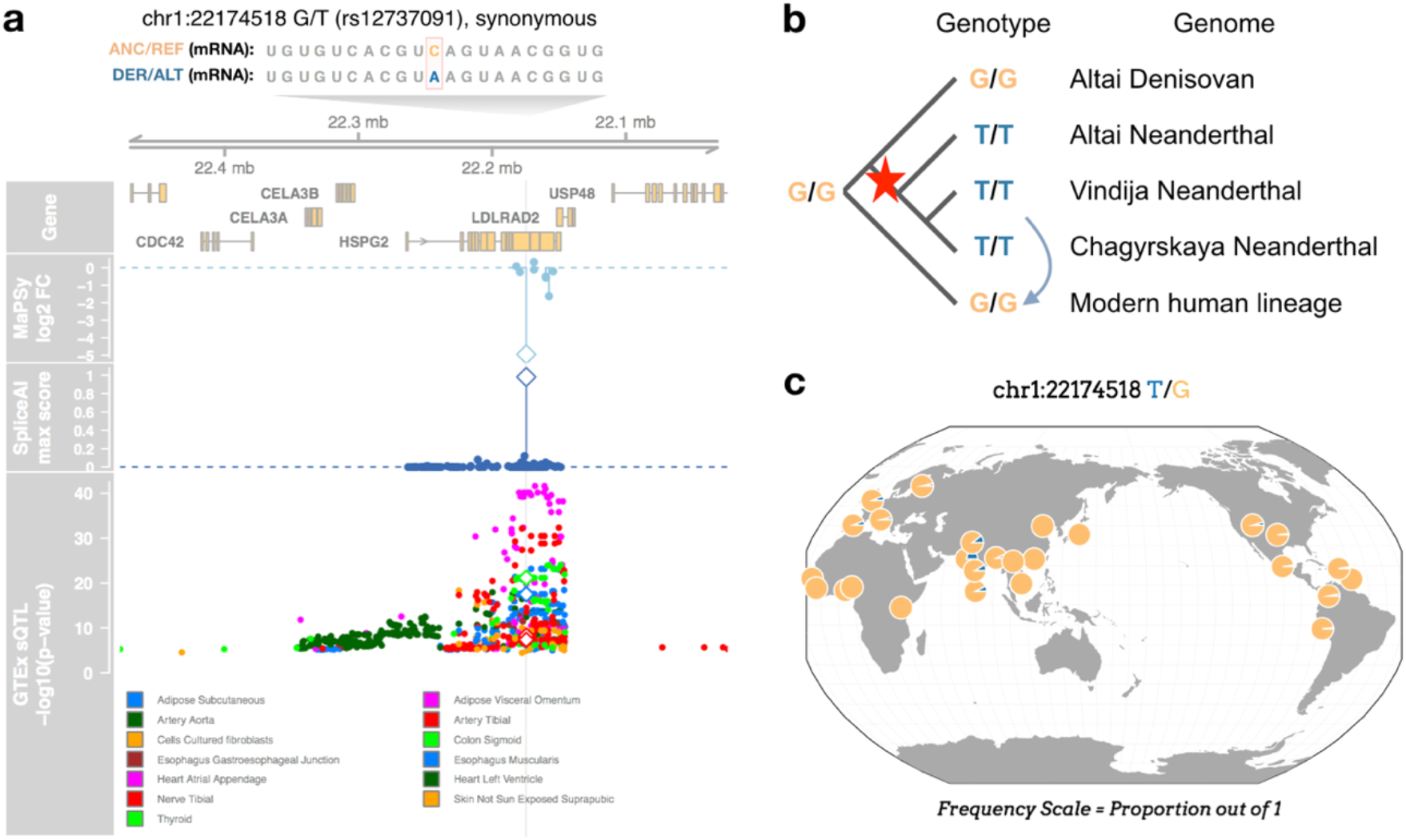
Neanderthal-specific and introgressed synonymous ESM (rs12737091) in *HSPG2*. The gene *HSPG2* encodes the ubiquitous extracellular matrix protein perlecan. **(a)** Genome tracks showing MaPSy, SpliceAI, and GTEx sQTL results for rs12737091 (diamonds) and nearby introgressed variants. **(b)** Comparisons between genotypes in four high-coverage archaic genomes and the consensus modern human genotype (red star represents origin of variant, light blue arrow represents introgression event). **(c)** 1KGP allele frequencies across 26 populations from the Geography of Genetic Variants Browser.

The previous case-studies of top lineage-specific ESMs were all introgressed. We further sought to identify a subset of high-confidence lineage-specific splicing variants that are private to either modern or archaic humans. We included all variants identified as MaPSy ESMs with a SpliceAI max raw score ≥ 0.1 that are not in the introgressed variant set and are not polymorphic in either 1KGP or gnomAD genomes. This resulted in 36 high-confidence private splicing variants, of which 51 were Denisovan-specific, 22 were Neanderthal-specific, 3 were archaic-specific, and none were modern-specific (Additional file 5: Table S4). Using more stringent splicing variant criteria of MaPSy strong ESMs with a SpliceAI max raw score ≥ 0.2 resulted in 9 private splicing variants, of which 8 were Denisovan-specific and 1 was Neanderthal-specific (Fig. S14). Variants in the more stringent set include (i) the Denisovan-specific splice region variant chr8:24342791 G/A in the gene *ADAM7*, which is important for sperm maturation and predicted by Combined Annotation Dependent Depletion to be deleterious (CADD score = 32) (82, 83); (ii) a Denisovan-specific splice region variant chrX:65247366 T/C in the gene *VSIG4*, which is a macrophage complement receptor and T-cell negative regulator (CADD SCORE = 19.8) (84, 85), and which lies in the middle of a previously reported introgression desert (35); and (iii) a Neanderthal-specific missense variant chr2:26702482 T/C in the gene *OTOF* which is involved in non-syndromic recessive deafness (CADD score = 11.1) (86).

### MaPSy identifies candidate trait-associated causal splicing variants

We sought to identify potential phenotypic impacts of causal splicing variants by exploring whether they were associated with trait variation. Most variants identified by genome-wide association studies (GWAS) are not causal due to LD. Statistical fine-mapping methods identify likely causal variants within a given phenotype-associated genomic region (87, 88). We intersected MaPSy ESMs with variants with posterior inclusion probability (PIP) ≥ 0.1 from a recent fine-mapping study across 94traits in the UK Biobank (UKB) and 79 traits in BioBank Japan (BBJ) (89). This identified seven variants across and eight phenotypes with evidence of both splicing and phenotypic impacts (Additional file 6: Table S5). However, 6 of these ESMs were also missense variants, suggesting alternative mechanisms of influencing phenotype. We identified one synonymous introgressed ESM, rs11045818 in the gene *SLCO1B1*, which has a moderately high PIP=0.626 for total bilirubin (Fig. S15a). *SLOC1B1* is primarily expressed in the liver and encodes the protein organic anion transporting polypeptide 1B1 (OATP1B1), which transports bilirubin and other compounds into the liver (90). The variant has a MaPSy functional score of −2.05 and a SpliceAI acceptor loss score of 0.09. The derived allele is fixed across all three Neanderthal genomes, and it narrowly misses our criteria for being identified as a Neanderthal-specific variant (1KGP and gnomAD AFR AFs are slightly above 1%) (Fig. S15b,c). A search for RBP-binding motifs using the ATtRACT database (76) suggests the G>A mRNA change introduces potential binding sites for splicing factors SRSF2 and NOVA1/NOVA2 (Fig. S15d), though only SRSF2 is highly expressed in liver tissue (91). The variant resides in a credible set with two intronic variants, rs17329885 (PIP=0.177) and rs11045820 (PIP=0.197), on the same introgressed haplotype. While rs17329885 is not predicted by SpliceAI to affect splicing, rs11045820 is predicted to have a potential small effect on splicing (acceptor loss score of 0.08), and thus may be a potential candidate causal variant in the credible set. Alternative splicing of *SLCO1B1* is not reported in GTEx V8 adult liver tissue (91), but has been reported in pediatric liver tissue (92) and in GTEx V9 long-read sequenced novel transcripts (93). We also intersected variants with SpliceAI max raw score ≥ 0.1 in the human evolution variant sets with fine-mapped variants with PIP ≥ 0.1. This identified an additional eight intronic introgressed variants, including one splice region variant, across seven distinct genes and 11 phenotypes (Fig. S16 and Additional file 7:Table S6). This included one adaptively introgressed intronic variant, rs17730088 in the keratin gene *KRT71* fine-mapped to balding type 4 (PIP=0.147), and two high PIP (≥0.5) introgressed intronic variants, rs2272783 in *FECH* fine-mapped to mean corpus hemoglobin (PIP=0.989) and rs8192327 in *SFTPC* fine-mapped to FEV/FVC ratio (PIP=0.875).

## Discussion

Key to our understanding of differences between modern and archaic humans is an appraisal of the functional effects of non-coding variants that existed in the evolutionary past, but which are no longer extant in present-day populations (5, 6, 41, 94, 95). Most genomic approaches developed so far have focused on transcriptional (31–33, 39, 40, 43) rather than post-transcriptional variant effects (but see (41) and (32)), are limited to studying introgressed variants associated with phenotypes in well-sampled modern human populations (4, 24, 35, 36, 38), and do not distinguish causal variants from linked variants. Recently, massively parallel reporter assays (MPRAs) have been used to address the latter two limitations, as shown in recent applications to detecting modern-specific and adaptively introgressed variants that affect cis-regulatory elements (50–53).

In this study, we applied a Massively Parallel Splicing Assay (MaPSy) (60) to detect 969 exonic splicing mutations (ESMs) out of a total of 5,224 tested variants, including nearly all modern-specific, archaic-specific, Neanderthal-specific, archaically introgressed, and adaptively introgressed variants that could fit in the assay (Fig. 1). We found MaPSy to be sensitive at detecting ESMs affecting both splice sites and exonic splicing regulatory elements (Fig. 2a, Fig. S5). To better understand the evolutionary landscape of splicing effects across the different variant sets in human evolution, we used a combination of insights from MaPSy functional scores, SpliceAI predicted splicing effects, and enrichment of GTEx sQTLs. We found that modern humans are depleted in loss of exon variants compared to the Neanderthal lineage (Fig. 2b–f), which is consistent with stronger purifying selection acting in the modern lineage due to their larger long-term effective population sizes (2, 16, 17). We next found that adaptively introgressed variants were enriched in moderate effect splicing variants relative to all introgressed variants (Fig. 3a). Analysis of enrichment or depletion of GTEx sQTLs and SpliceAI scores (relative to non-introgressed control variants with matching AF and LD) showed that this was likely due to both increased purifying selection acting against introgressed variants that result in loss of exon inclusion, and positive selection in favor of moderate effect, but not weak or strong effect, splicing variants that altered exon inclusion frequencies in modern humans (both positively and negatively) relative to the hominin ancestor (Fig. 3b–h, Fig. S10). This was somewhat unexpected, and suggests the hypothesis that adaptive introgression may favor functional alleles with intermediate effect sizes. Weak to moderate splicing variants are more likely to contribute to tissue-specific alternative splicing (72). Selection coefficients may be more likely to be positive if variants only disrupt splicing in some tissues, and more likely to be negative if variants strongly disrupt splicing throughout the body.

Finally, we provide evidence that MaPSy can identify candidate causal splicing variants. As an illustrative example, we characterized an adaptively introgressed ESM, rs5743566, in the innate immunity gene *TLR1* (Fig. 4). The *TLR6/TLR1/TLR10* locus is a hotspot for repeated archaic introgression due to an important role in recognizing pathogens (29). Introgressed haplotypes at this locus have been shown to affect both *TLR1* gene expression and splicing (22, 29, 38–41). The MaPSy-identified variant, rs5743566, lies in in an internal exon in the 5’ UTR of *TLR1* away from the splice regions. It is thus a seemingly unlikely candidate for splicing effect annotation. However, MaPSy functional scores, SpliceAI scores, predicted changes in RBP binding motifs, and predicted changes in exonic splicing enhancer activity, all supported the conclusion that rs5743566 is a causal splicing variant with a moderate negative effect on *TLR1* exon 2 inclusion (Fig. 3). Moreover, GTEx identifies rs5743566 as a significant sQTL in seven out of 49 tissues, which could be explained by tissue-specific alternative splicing (91). Interestingly, Jagoda et al. (53) recently found another putatively causal variant, rs73236616, at this adaptively introgressed locus in a cis-regulatory element that impacts gene expression not in *TLR1, TLR6*, or *TLR10*, but the neighboring *FAM114A1* gene. Thus, multiple causal variants affecting different genes may be found on the same adaptively introgressed haplotype, any of which could be a target of positive selection. We further showed that MaPSy can identify causal splicing variants that are fixed or nearly fixed on the modern, archaic, and Neanderthal lineages (Fig. 5 and S12-14), such as the derived Neanderthal-specific ESM, rs12737091, in the gene *HSPG2* that encodes the ubiquitous extracellular matrix protein perlecan, and the derived modern-specific ESM, rs77724956, in the gene *FAAH* that is also an example of Neanderthal introgression reintroducing an ancestral functional allele back into modern populations at low frequencies (Fig. S13) (51). This also includes a set of high confidence splicing variants private to Neanderthals or Denisovans that are not found in 1KGP or gnomAD genomes, and would be unlikely to have been discovered even with deep resampling of transcriptomes across diverse modern human populations (Fig. S14). These include variants that strongly disrupt splicing and are predicted to be highly deleterious by CADD scores in genes related to sperm maturation, *ADAM7*, and immune response, *VSIG4*. Finally, we identified MaPSy and SpliceAI-identified candidate splicing variants that are also candidate causal variants for trait-associations in the UK Biobank and BioBank Japan (89), including variants underlying associations with total bilirubin, balding type 4, mean corpus hemoglobin, and FEV/FVC ratio (Fig. S15-S16).

There are notable limitations of MaPSy that apply to our study. While we validated a half exon approach to testing exonic variants near the 3’ ss of longer exons (Fig. S2), there are still many exonic variants that cannot be tested using MaPSy. Intronic variants, which include those that disrupt canonical splice signals, such as the 3’ ss, 5’ ss, polypyrimidine tract, and branchpoint signal, were not tested using MaPSy. These could be tested using newer splicing assay designs based on barcoding or fluorescence-activated cell sorting (63, 64). The minigenes used by MaPSy can only include a small part of endogenous sequence, and thus lack important contextual sequences that are further away and that also help determine splicing (72), such as potential competing splice sites of neighboring exons (96). Another limitation is that we only transfected a single cell type, HEK 293T. This is less of a problem for MaPSy than other MPRAs, because splicing variation is shared across tissues more than expression variation (91, 97). The notable exception is brain-related tissues, which have a distinct splicing program compared to other tissues (97), and express tissue-specific splicing factors that regulate alternative splicing, such as *NOVA1* (98). Neuronal cell lines and primary cells would thus be a high priority for future MaPSy experiments. As well, MaPSy only evaluates individual variants with cis-effects on splicing. Trans-effects on splicing are also important, and notably, all modern humans carry a fixed derived mutation in *NOVA1* that has been shown to alter transcriptomes and neurodevelopment compared to the ancestral allele found in archaic humans (46). It is also conceivable that multiple variants in cis could have synergistic effects on splicing (61, 99); this aspect of splicing is also not interrogated by MaPSy.

Our use of MaPSy provides novel insights into role of natural selection acting on splicing variation in hominin evolution, and enabled the identification of causal splicing variants, not limited to those in well-sampled present-day populations. Approaches such as MaPSy thus open the door to functionally characterizing variants found in diverse, understudied, and extinct populations. MaPSy is but one of many recently developed multiplexed assays of variant effect (100). As these assays and other emergent methods for interpreting variant-to-function improve, such as gene editing and machine learning models, we will increasingly be able to apply them to answer questions about the origins of our species’ unique molecular and complex phenotypes.

## Materials and Methods

### Selection of variants in human evolution variant sets

Individual variant sets were described in the main text in the section “Defining human evolution variant sets used in the MaPSy experiment” and also in detail in Fig. S3. All analyses used the hg19 reference. For lineage-specific variants, we consider the cases where the human reference (REF) allele is either the ancestral (ANC) or derived (DER) allele separately, based on inferred ANC alleles from Ensembl (70) from ftp://ftp.ensembl.org/pub/release75/fasta/ancestral_alleles/homo_sapiens_ancestor_GRCh37_e71.tar.bz2, and ignoring sites with ambiguous ANC alleles. These were then filtered based on 1KGP phase 3 global or African AF from http://hgdownload.cse.ucsc.edu/gbdb/hg19/1000Genomes/phase3/ and gnomAD (v2.1.1 genomes) global or African AF from https://gnomad.broadinstitute.org/downloads (68). Finally, these were then filtered by archaic allele counts in the four high-coverage archaic genomes from http://cdna.eva.mpg.de/neandertal/, and by the recommended filters for each archaic genome. 1KGP and gnomAD AF information could be missing due to either a true absence of the ALT allele in the population or because they were not genotyped. If AF is missing in one of 1KGP or gnomAD but not the other, we assumed that it was not genotyped in the dataset where it is missing. If AF is missing in both of 1KGP or gnomAD, we assumed that it was not genotyped or truly absent depending on whether it was found at low or high AF across the four high-coverage archaic genomes. Introgressed variants were downloaded from https://akeylab.princeton.edu/downloads.html (8) and https://data.mendeley.com/datasets/y7hyt83vxr/1 (11). Adaptively introgressed variants were downloaded from https://doi.org/10.1016/j.cub.2016.10.041 (Supplementary Table S1 of (22)) and Racimo https://doi.org/10.1093/molbev/msw216 (Supplementary Table S3 of (23)). This resulted in 2,034,092 total variants across variant sets (Additional file 2: Table S1). Variant set intersections were plotted using UpSetR (101).

### Selection of variants used in MaPSy experiment

GENCODE v32 basic transcripts with liftOver to hg19 were downloaded from https://ftp.ebi.ac.uk/pub/databases/gencode/Gencode_human/release_32/gencode.v32.basic.annotation.gff3.gz. A single canonical transcript for each gene was retained based on APPRIS annotations (selection criteria in descending order of priority: appris_principal_1 through 5, appris_alternate_1 through 2, and then longest transcript) (102). Variants in the human evolution variant sets were then overlapped with internal exons in the transcript set. For short exons (10 ≤ length ≤ 120 bp), all exonic variants were retained. For long exons (121 ≤ length ≤ 500), only exonic variants within the first 90 bp from the 3’ ss were retained. This resulted in an initial set of 6,376 variant-exon pairs for the MaPSy experiment (Additional file 3: Table S2). This includes additional variant-exon pairs found only in two earlier studies (1, 2) of lineage-specific variants that were later dropped and not retained in the final analysis.

### MaPSy oligo library design and experiment

See additional text in SI Appendix.

### SpliceAI score annotations

Precomputed genome-wide raw SpliceAI scores (v1.3) were downloaded from https://basespace.illumina.com/s/otSPW8hnhaZR (72). SpliceAI max scores were defined as the max of raw SpliceAI donor gain, donor loss, acceptor gain, and acceptor loss scores. Variants that overlap multiple genes were assigned to the one with highest SpliceAI max raw score. Variants were binned by SpliceAI max scores into no splicing (score of 0), weak (between 0.01 and 0.19), moderate (between 0.2 and 0.49), and strong (0.5 to 1) effect categories. Pairwise differences in proportions were tested using Fisher’s Exact Test. Enrichment analyses for each SpliceAI effect category were based on AF-matched non-introgressed controls. SpliceAI scores are available in Additional file 2: Table S1.

### Hexamer scores of exonic splicing regulatory activity

Hexamer-based scores for exonic splicing regulatory based on high-throughput minigene experiments were downloaded from two previous studies. Enrichment index (EI) hexamer scores were downloaded from https://genome.cshlp.org/content/21/8/1360/suppl/DC1 (Supplementary Table 1 of (73)). Exonic alternative 3’ ss (exonic A3SS) and 5’ ss (exonic A5SS) hexamer scores were downloaded from https://github.com/Alex-Rosenberg/cell-2015/blob/master/ipython.notebooks/Cell2015_N4_Motif_Effect_Sizes.ipynb (74). Delta EI scores for a variant in a gene were calculated as the mean of six overlapping hexamer EI scores on the coding strand for the DER sequence minus that of the ANC sequence. Delta exonic A3SS and Delta exonic A5SS scores were defined similarly. Hexamer score annotations for MaPSy variants are available in Additional file 3: Table S2.

### Ensembl VEP annotations

The Ensembl Variant Effect Predictor (VEP) command line tool (v101.0) (103) was used to annotate variants with VEP consequences using the parameters: --cache --pick --assembly GRCh37. Analysis of VEP proportions by MaPSy functional score bins was restricted to exonic variants with splice region, stop gain, missense, synonymous, 5’ UTR, and 3’ UTR VEP annotations. Enrichment analyses of VEP annotations were based on AF-matched non-introgressed controls. VEP annotations are available in Additional file 3: Table S2.

### GTEx sQTL and eQTL annotations

GTEx (V8) sQTLs (eQTLs) significant at FDR < 0.05 for 49 tissues defined on GRCh38 were downloaded from https://gtexportal.org/home/datasets (91). These were then converted to hg19 coordinates using CrossMap (v0.5.2) (104). Variants involved in a sQTL (eQTL) association in at least one tissue were said to overlap a sQTL (eQTL) (Additional file 10: Table S9 and 11: Table S10).

### Defining non-introgressed matched controls

Enrichment analyses were conducted using the subset of 1000 Genomes Project (1KGP) phase 3 autosomal single nucleotide polymorphisms (SNPs) found in the European superpopulation with a minor allele frequency (MAF) of at least 0.01. This was because the vast majority of individuals in GTEx are of European descent, and GTEx QTL results are only available for SNPs with MAF > 0.01. 1KGP SNPs on chromosomes 1-22 in hg19 were downloaded from http://hgdownload.cse.ucsc.edu/gbdb/hg19/1000Genomes/phase3/ (67). 1KGP SNPs were initially binned by their European allele frequency (EUR AF) into 50 AF bins (allele counts of 1 through 10 plus 40 equal sized bins for allele counts > 10), and by a univariate summary of LD in Europeans (EUR LD, defined for a given SNP as the number of SNPs with European minor allele frequency (MAF) > 0.01 within 1 Mb with r^2^ > 0.2) into 20 equal sized LD bins. LD between pairs of 1KGP SNPs was calculated using vcftools (v0.1.16) (105) using the parameters: --hap-r2 --maf 0.01 --ld-window-bp 1000000 --min-r2 0.2 --max-missing 1. Finally, the set of non-introgressed controls was defined as the subset of 1KGP autosomal SNPs with European MAF > 0.01 not already found in the introgressed variant set (Additional file 12: Table S11).

### Enrichment analyses relative to matched non-introgressed controls

We describe the enrichment analyses for GTEx sQTLs among introgressed variants, but all enrichment analyses are conducted similarly. For introgressed variants in a AFxLD bin, we sample with replacement an equal number of non-introgressed variants belonging to the same AFxLD bin. We then calculate the log_2_ odds-ratio that the introgressed variants overlaps a sQTL in at least one tissue compared to the set of matched non-introgressed controls. We do this for 1,000 randomly sampled sets of matched non-introgressed controls to quantify the variation in log_2_ odds-ratio estimates. The p-value for the two-tailed test of the null hypothesis that the log_2_ odds-ratio equals 0 is calculated as 2*min(fraction of estimates above 0, fraction of estimates below 0). Finally, to test for a difference between the log_2_ odds-ratio estimates for introgressed versus adaptively introgressed variants, we calculate a two-tailed p-value in the same way but for the 1,000 differences between their log_2_ odds-ratios.

VEP and SpliceAI annotations do not depend on LD, whereas GTEx QTLs are highly dependent on LD. Enrichment analyses for VEP annotations were done using 100 randomly sets of AF-matched non-introgressed controls, for GTEx eQTLs and sQTLs using 1,000 randomly sampled sets of AF and LD matched non-introgressed controls, and for SpliceAI categories using 1,000 randomly sampled sets of AF matched non-introgressed controls.

Because the background sets used are the set of exonic variants included in the MaPSy experiment, and the set of variants that could be scored by SpliceAI within 50 bp of an exon, this analysis controls for the known enrichment of genes overlapping adaptive introgression.

### Visualizations for causal splicing variant case studies

Genome tracks for MaPSy functional scores, SpliceAI scores, and GTEx sQTL p-values were visualized using Gviz (v1.32.0) (106). SpliceAI scores were only shown for other variants that share a variant set with the target variant. GTEx results were shown only for other variants associated with the same gene as the target variant. 1KGP population allele frequencies were downloaded from the Geography of Genetic Variants Browser (https://popgen.uchicago.edu/ggv/) (107). The ATtRACT RBP motif database was used to search the DER or ANC 11-mer sequence window centered on a target variant for RBP binding motifs of at least 6 bp in length (76). eQTLGen Consortium whole blood eQTLs are from https://www.eqtlgen.org/cis-eqtls.html (81).

### Overlap with UK Biobank and BioBank Japan fine-mapped trait-associated variants

Fine-mapping results for 94 complex traits in the UK Biobank and BioBank Japan were downloaded from https://www.finucanelab.org/data and https://pheweb.jp/ (89). Fine-mapped variants with PIP ≥ 0.1 were overlapped with either MaPSy ESMs or variants with SpliceAI max raw score ≥ 0.1 taking the higher of FINEMAP or SuSiE PIP scores (Additional file 6: Table S5 and 7: Table S6).

### Availability of data and materials

Supplementary figures S1-S16 are available in Additional file 1. The datasets supporting the conclusions of this article are available in Additional files 2-12 deposited in Zenodo (https://doi.org/10.5281/zenodo.7225467). MaPSy sequencing data are deposited in Gene Expression Omnibus (accession GSE201856). Custom scripts for reproducing analyses, tables, and figures are available on GitHub (https://github.com/stephenrong/archaic-splicing). Large files for running custom scripts are available in Zenodo (https://doi.org/10.5281/zenodo.7225467).

## Acknowledgments

We would like to thank the Fairbrother Lab, Joaquin C.B. Nunez, and Emilia Huerta-Sanchez for discussions and input. This work was supported by the National Institutes of Health (R01 GM127472 to W.G.F.), and the Brown-MBL IGERT in Reverse Ecology (NSF 0966060). S.R. was supported by a NSF Graduate Research Fellowship (NSF 1644760).

## Author Contributions

S.R., W.G.F, C.R.N., and B.J.E designed the study. C.R.N. performed the MaPSy experiment. S.R. designed oligos for the MaPSy experiment, analyzed the MaPSy data, and conducted the analyses in the paper. S.M, I.C.M., and M.M annotated variants found in a pilot MaPSy experiment. S.R. and W.G.F. wrote the paper with contributions from all authors.

## Competing Interest Statement

W.F.G. is on the scientific advisory board of Remix Pharmaceutical and a founder of WALAH Scientific.

## Notes

https://doi.org/10.5281/zenodo.7225467

https://github.com/stephenrong/archaic-splicing/

https://www.ncbi.nlm.nih.gov/geo/query/acc.cgi?acc=GSE201856

### Summary of Updates

Updated Zenodo DOI in manuscript and updated author institutions and department affiliations.

