## Supplementary Material for "Large scale functional screen identifies genetic variants with splicing effects in modern and archaic humans"

**Supporting Information Text**

**Methods**

*MaPSy for variants in both short and long exons*

A limitation of previous MaPSy experiments is that they could only characterize ESMs in short exons (1). Because MaPSy relies on 230 nt oligo synthesis, and each oligo contains an exon plus flanking intronic sequence, only SNVs in the 42% of canonical transcript exons smaller than 120 bp are typically assayed. However, here, we extended the capabilities of MaPSy by showing that we can reliably use a “half exon” construct to characterize ESMs near the 3’ splice site (3’ ss) of longer exons (> 120 bp). Each half exon construct contains a truncation of a long exon to its first 100 bp with a SNV in the first 90 bp plus 55 bp of upstream intronic sequence containing the 3’ ss (Fig. S2a). To complete the exon, we ligated a shared sequence containing a 5’ splice site (5’ ss) with 20 bp of upstream exonic and 15 bp of downstream intronic sequence. This is then incorporated into the minigene experiment with other details following previous work (1). To validate this approach, we created half exon constructs for a sample of short exons that could be compared with their full exon counterparts, and tried two different 5’ ss-containing common sequences labeled 3A and 3B. Across pairwise comparison between constructs containing the same variant, we found strong agreement in their MaPSy $\log_2$ FC estimates (*r* between 0.79-0.83) (Fig. S2b-e). We performed a similar experiment for variants in the exon end containing the 5’ ss, but found only moderate agreement in pairwise comparisons (*r* between 0.42-0.78) (Fig. S2f-i). This is likely due to the inclusion of a much longer shared sequence, a much shorter endogenous sequence, and absences of the endogenous 3’ ss, polypyrimidine tract, and branchpoint signal. For this reason, only the full exon constructs for short exons (between 10 and 120 bp) and 3A half exon constructs for long exons (between 121 and 500 bp) were analyzed. This covers an improved 90% of all internal exons in canonical transcripts, excluding only the very shortest and longest exons in the genome, and nearly doubles the number of SNVs that could be assayed. MaPSy functional scores for matched validation constructs are available in Additional file 8: Table S7.

*Oligonucleotide pool design and synthesis*

We designed oligos corresponding to either the DER or ANC allele of each variant. Each 230 nt oligo contains in 5’ to 3’ order a 25 nt forward primer, 190 nt of variable sequence, and a 15 nt reverse primer. For short exons (10 ≤ length ≤ 120 bp), the 190 nt variable sequence contains at least 55 nt of the upstream intronic sequence, the endogenous exon containing the variant allele of interest, and 15 nt of the downstream intronic sequence. For long exons (121 ≤ length ≤ 500), we used the 3A half exon construct for variants in the first 90 nt of the exon near the 3’ ss. The 190 nt variable sequence contains 55 nt of upstream intronic sequence, the first 100 nt of the endogenous exon containing the variant allele of interest (including a 10 nt buffer region at the end), and the common 3A sequence containing a 5’ ss with 20 nt of the upstream exon and 15 nt of the downstream intron (common 3A, Additional file 9: Table S8). Very long exons (> 500 nt) were not included in the analysis. The oligo sequences were grouped into separate oligo sublibraries (primer pair 01-08, Additional file 9: Table S8). The oligo library pool was synthesized by Agilent Technologies and used to generate minigene reporters.

*Oligo design for half exon validation MaPSy experiment*

In a pilot MaPSy experiment, we used a subset of the human evolution variants defined in this study along with positive control variants identified as ESMs in an earlier MaPSy study (1) to validate the half exon construct approach. In addition to short full exon constructs, we designed short half exon constructs for either variants near the 3’ ss (using common 3A or 3B sequences) or the 5’ ss (using common 5A or 5B sequences). For the short half 3A and 3B constructs corresponding to exons of length *L*, the 190 nt variable sequence contains at least 55 nt of the upstream endogenous intron, *L*–10 nt of the endogenous exon containing the variant allele of interest in the first *L*–20 nt, and either the 3A or 3B common sequence containing 20 nt of upstream exon and 15 nt of downstream intron. For the short half 5A and 5B constructs, the 190 nt variable sequence contains either the 5A or 5B common sequence containing 55 nt of upstream intron and 20 nt of downstream exon, the last *L*–10 nt of the endogenous exon containing the variant allele of interest in the last *L*–20 nt, and 15 nt of downstream endogenous intron. Long half 3A, 3B, 5A, and 5B constructs are constructed similarly, using only the first or last 100 nt of the endogenous exon sequence.

*MaPSy minigene construction*

The MaPSy minigene reporter construct (Fig. S1a) includes from 5' to 3' order a cytomegalovirus (CMV) promoter (CMV promoter), an adenovirus pHMS91 exon (AD81) exon with part of its downstream intron (AD81 exon-intron), a variable 230 bp sequence from oligo synthesis (oligo sequence, plus an oligo extension of the reverse primer by 15 bp), exon 16 of the ACTIN1 gene with part of its upstream intron (ACTIN intron-exon), and a bGH polyA signal containing sequence (bGH polyA) (Additional file 9: Table S8). 5' upstream (CMV promoter + AD81 exon-intron) and 3' downstream (oligo extension + ACTIN intron-exon + bGH polyA) minigene fragments were extended to include sequence unique to each of the oligo sublibraries using extension PCR, and full minigenes were subsequently concatenated using overlapping PCR. The minigene libraries were then pooled together in equimolar amounts, resulting in an input library with all minigene reporter constructs.

*MaPSy transfection and input and output library sequencing*

The input library was transfected into human embryonic kidney HEK 293T cells obtained from the American Type Culture Collection (ATCC CRL-316) in three cell culture replicates using Lipofectamine 3000 (Invitrogen) in a 6-well plate. HEK 293T is not listed in the ICLAC Register of Misidentified Cell Lines (v10) and was confirmed to be mycoplasma free in previous passage. Forty-eight hours after transfection, RNA was extracted using TRIzol (ThermoFisher) and DNase treated (Invitrogen). Random 9-mers were used to generate cDNA with SuperScript IV Reverse Transcriptase (Invitrogen) followed by 25 cycles of PCR amplification (GoTaq, Promega), resulting in output libraries of spliced species.

Primers were designed to amplify the region of the input and output library sequences immediately surrounding the variable region of interest (input and output primer pairs, Additional file 9: Table S8). For the input libraries, the primers matched AD81 and ACTIN1 intronic regions. For the output libraries, the primers matched exonic regions of AD81 and ACTIN1. Input libraries of minigene reporters and output libraries of spliced species were then sequenced on Illumina HiSeq 4000 (2x150 paired-end) with one lane for three input libraries (technical replicates), and a second lane for three output libraries (transfection replicates). Replicates were identified using NEXTflex DNA Barcodes (PerkinElmer).

*Alignment and QC of MaPSy reads*

STAR (v2.7.9a) (2) was used to perform end-to-end unspliced alignment of paired-end reads against custom reference genomes with either input minigene sequences or the expected output three exon sequences with up to 5 mismatches and no indels using the parameters: --alighIntronMax 1 --outFilterMismatchNmax 5 --scoreDelOpen -1000 --scoreInsOpen -1000 --alignEndsType EndToEnd --outFilterMultimapNmax 1. Samtools idxstats (v1.7) (3) was used to count the number of reads aligned to each allelic species in each input and output sequencing library. The resulting read counts were then joined to a table of variant-exon pairs. We performed quality control (QC) of variant-exon pairs by applying the following filters: DER and ANC read counts of at least 20 in all three input and all three output replicates; and at least one mapped DER or ANC read in at least one of the three output replicates. We also removed variants only found in two earlier studies of lineage-specific variants (4, 5). After QC, we retained 5,224 variant-exon pairs belonging to at least one of the variant sets for analysis (Additional file 3: Table S2).

*MaPSy functional scores from read counts*

The MaPSy functional score is defined as the $\log_2$ fold change (FC) of derived (DER) versus ancestral (ANC) read counts plus a pseudocount of 1 in the output library normalized by read counts plus a pseudocount of 1 in the input library, or $log_2\big((DER_o/ANC_o)/(DER_i)/(ANC_i)\big)$. To account for variability across replicates, MaPSy functional scores and p-values for the null hypothesis that $\log_2$ FC = 0 were estimated using the method mpralm (6), a variation of the limma-voom framework for differential expression in MPRAs (7, 8). We used the mpralm $\log_2$ FC estimate and moderated t-test p-values with FDR correction for multiple tests (9). We identified significant ESMs at FDR < 0.05 and |FC| > 1.5. Weak ESMs had |FC| between 1.5 and 3. Strong negative ESMs had FC < -3. Strong positive ESMs had FC > 3. Pairwise differences in proportions were tested using Fisher's Exact Test. MaPSy functional scores are available in Additional file 3: Table S2.

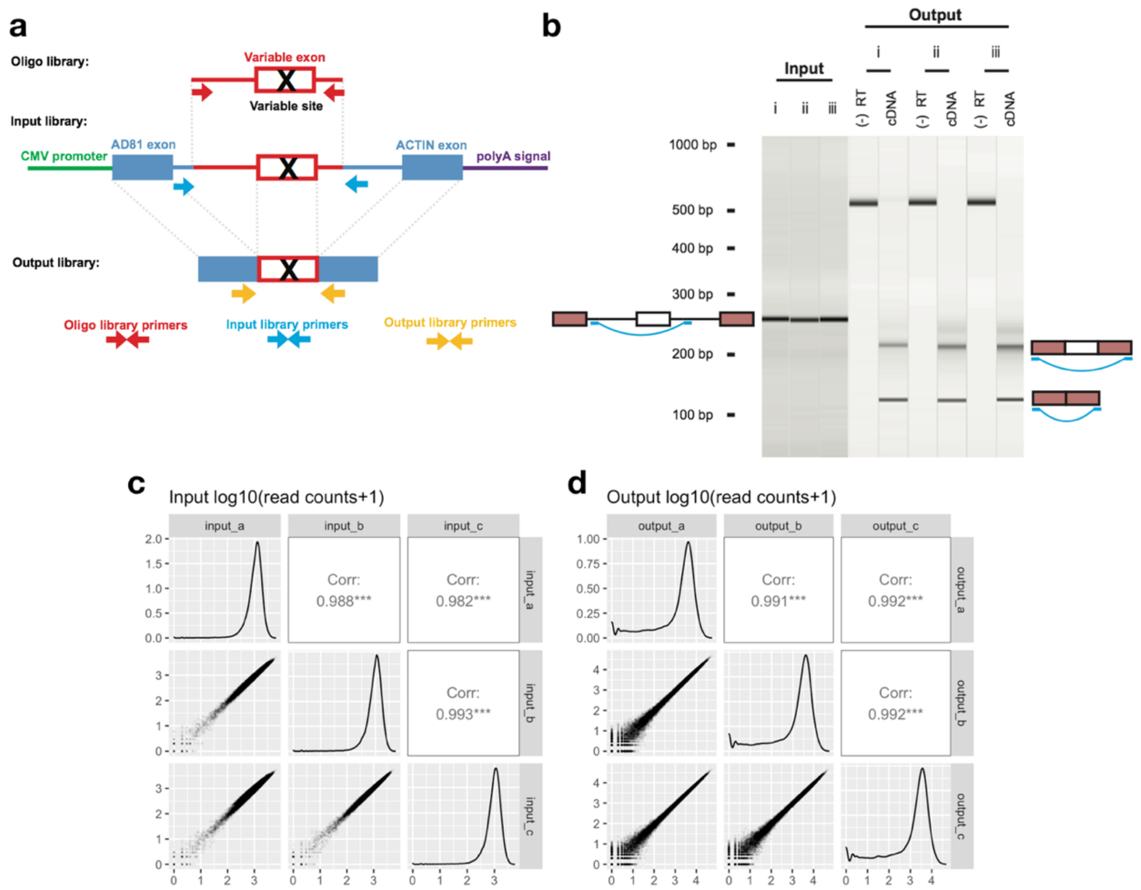


**Fig. S1. MaPSy minigene construction and reproducibility. (a)** Diagrams of MaPSy oligo library, input minigene constructs, and output spliced transcripts, and target locations of amplicon primer pairs. **(b)** RT PCR gel from the human evolution MaPSy experiment. Shown are bands from three replicates of input DNA library amplicons, output cDNA library amplicons in either the case where the middle exon is included or skipped, and minus RT controls. **(c)** Reproducibility of input DNA sequencing read counts. **(d)** Reproducibility of output cDNA sequencing read counts. Pearson's correlation coefficient between log10 read counts + 1 pseudocount.

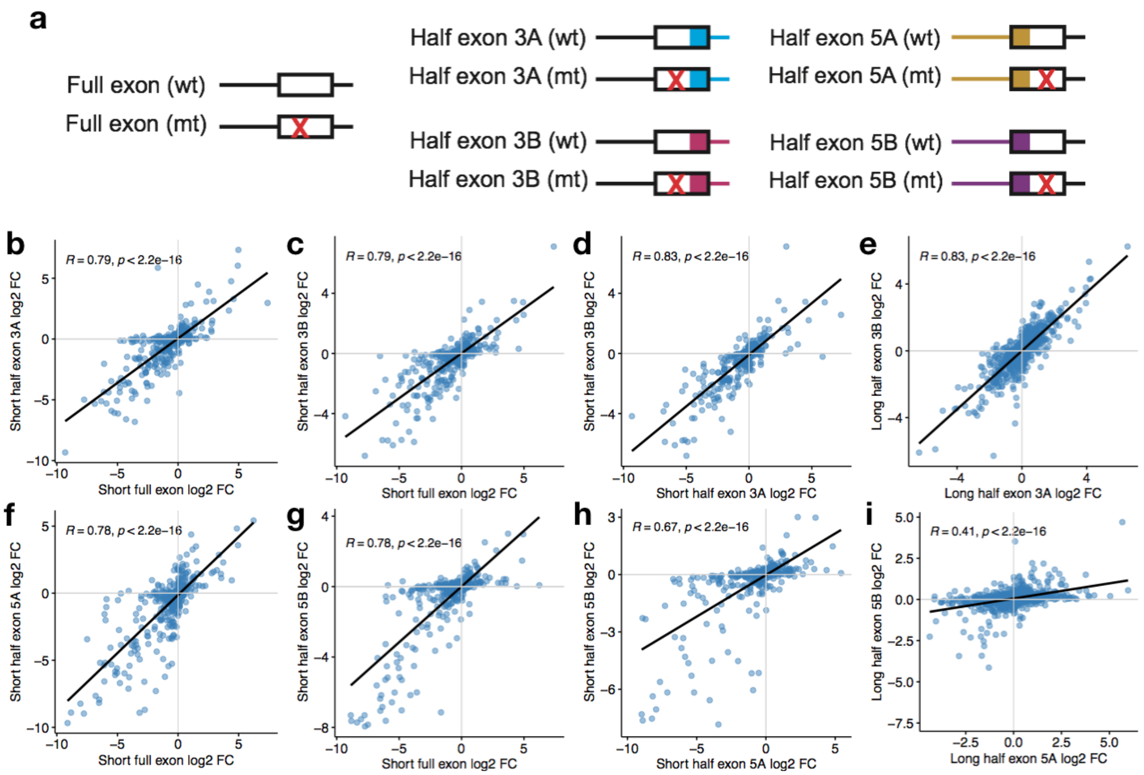


**Fig. S2. Validation of use of 3' ss half exon constructs in MaPSy. (a)** Diagrams of full exon constructs, two versions 3A and 3B of the 3' ss half exon constructs, and two versions 5A and 5B of the 5' ss half exon constructs used in the half exon validation MaPSy experiment. Short exons (≤ 100 bp) represented by full exons, 3A, 3B, 5A, and 5B half exons. Long exons (>100 bp) represented by 3A, 3B, 5A, and 5B half exons. Comparisons of MaPSy functional scores (log2 FC) between: **(b)** short full exons vs 3A half exons, **(c)** short full exons vs 3B half exons, **(d)** short 3A vs 3B half exons, **(e)** long 3A vs 3B half exons, **(f)** short full exons vs 5A half exons, **(g)** short full exons vs 5B half exons, **(h)** short 5A vs 5B half exons, and **(i)** long 5A vs 5B half exons.

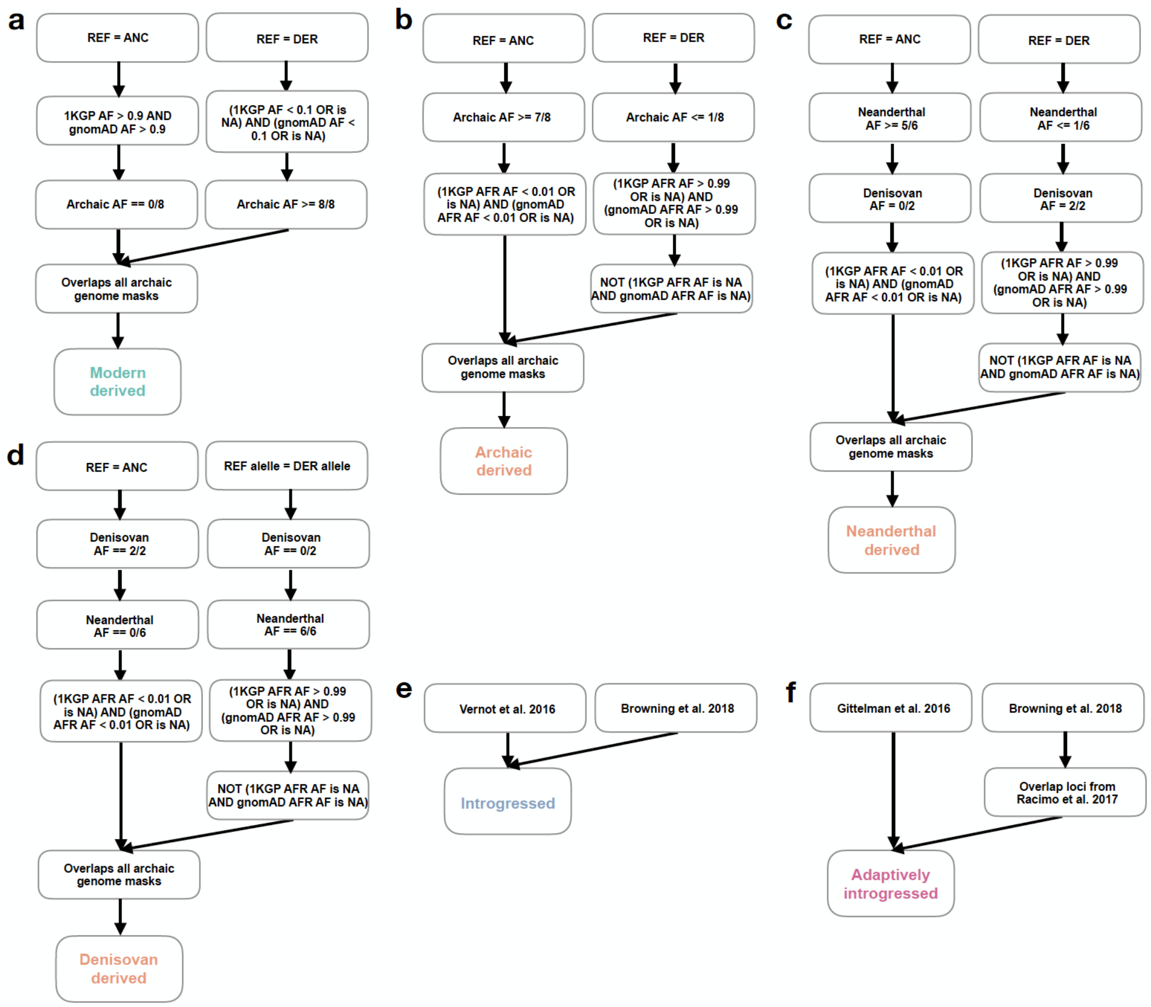


**Fig. S3. Definitions of the main variant sets used in this study.** Decision trees showing how single nucleotide variants were assigned to each variant set: **(a)** modern-specific, **(b)** archaic-specific, **(c)** Neanderthal-specific, **(d)** Denisovan-specific, **(e)** introgressed, and **(f)** adaptively introgressed.

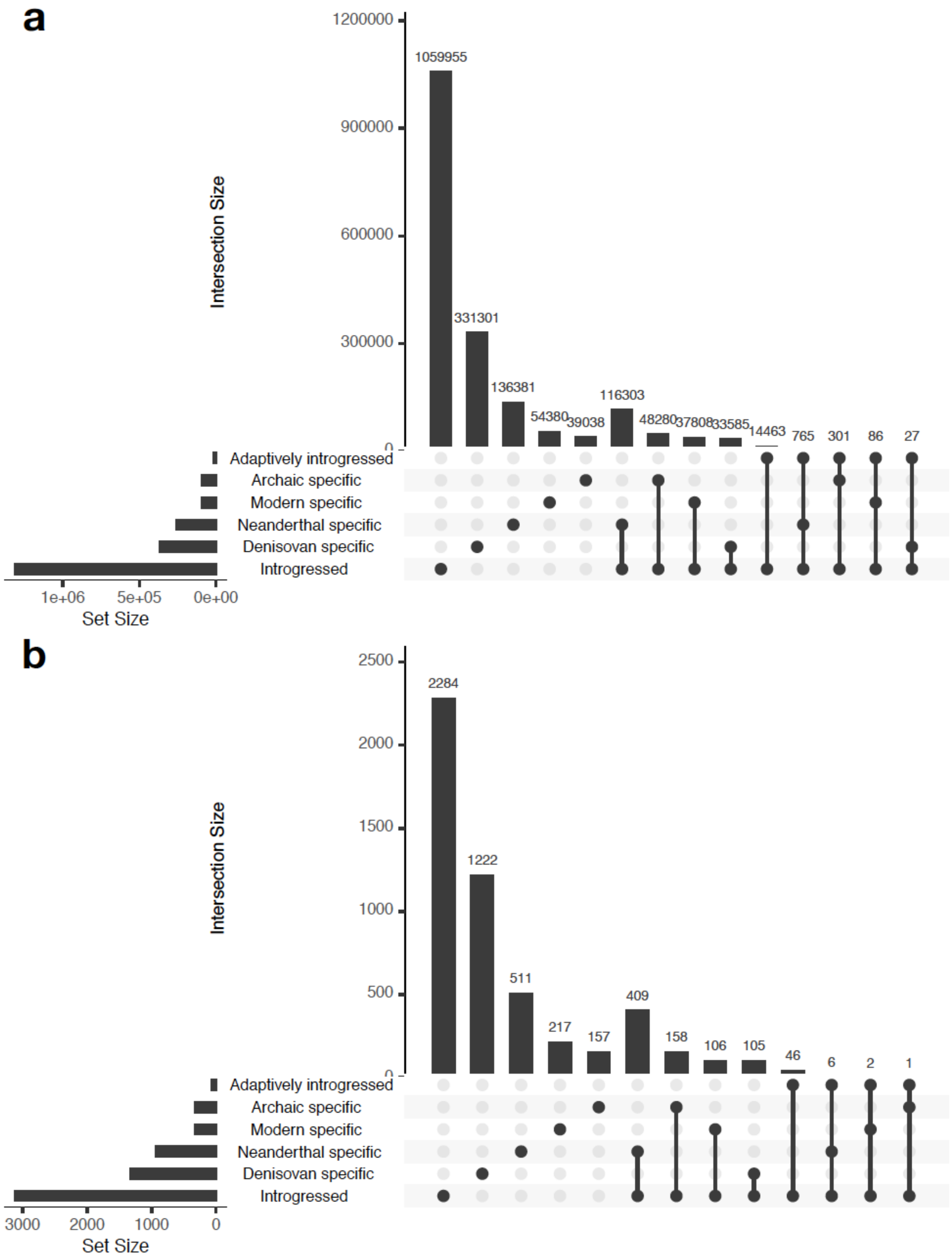


**Fig. S4. Overlaps between main variant sets in this study.** UpSet plots showing the number of variants in each nonempty intersection between variant sets for **(a)** all variants in the human genome, and **(b)** only the variants assayed using MaPSy.

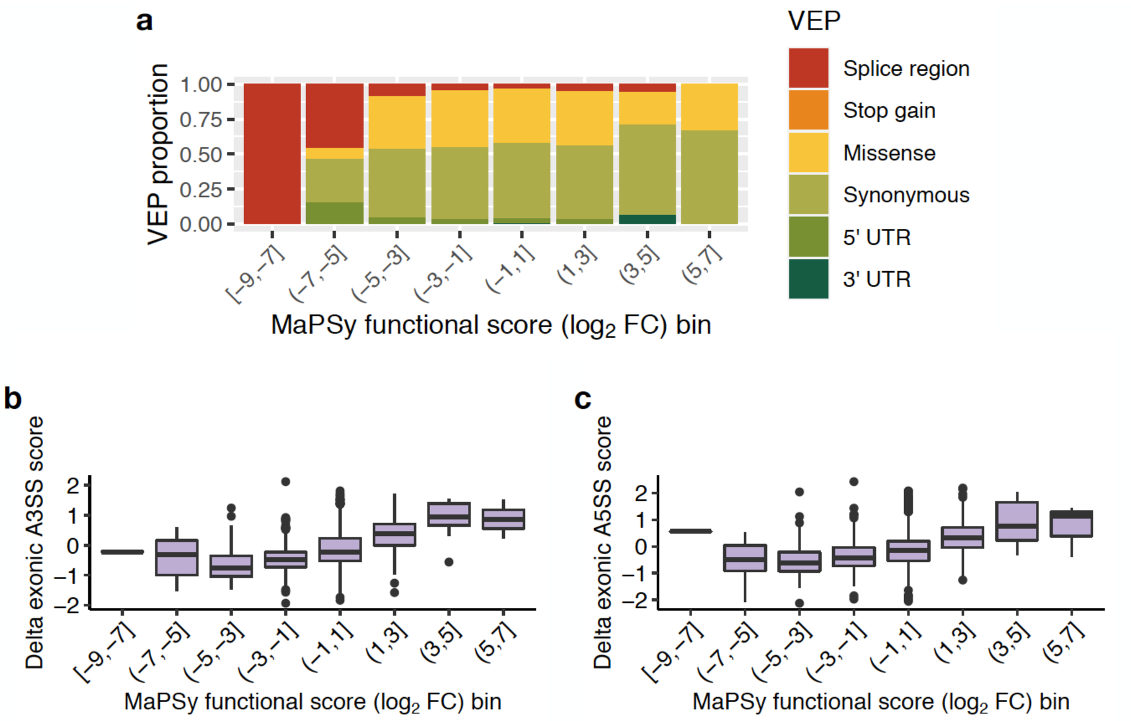


**Fig. S5. Relationship between MaPSy functional scores, VEP consequence, and alternative measures of change in exonic splicing enhancer activity.** **(a)** Ensembl Variant Effect Predictor (VEP) consequence proportions by MaPSy functional score (log2 FC) bins. Relationship between MaPSy functional score (log2 FC) bins and: **(b)** Delta exonic A3SS score, the change in the mean of overlapping hexamers scored empirically for their exonic splicing enhancer activity in the context of alternative 3’ ss usage, **(c)** Delta exonic A5SS score, defined similarly, but in the context of alternative 5’ ss usage.

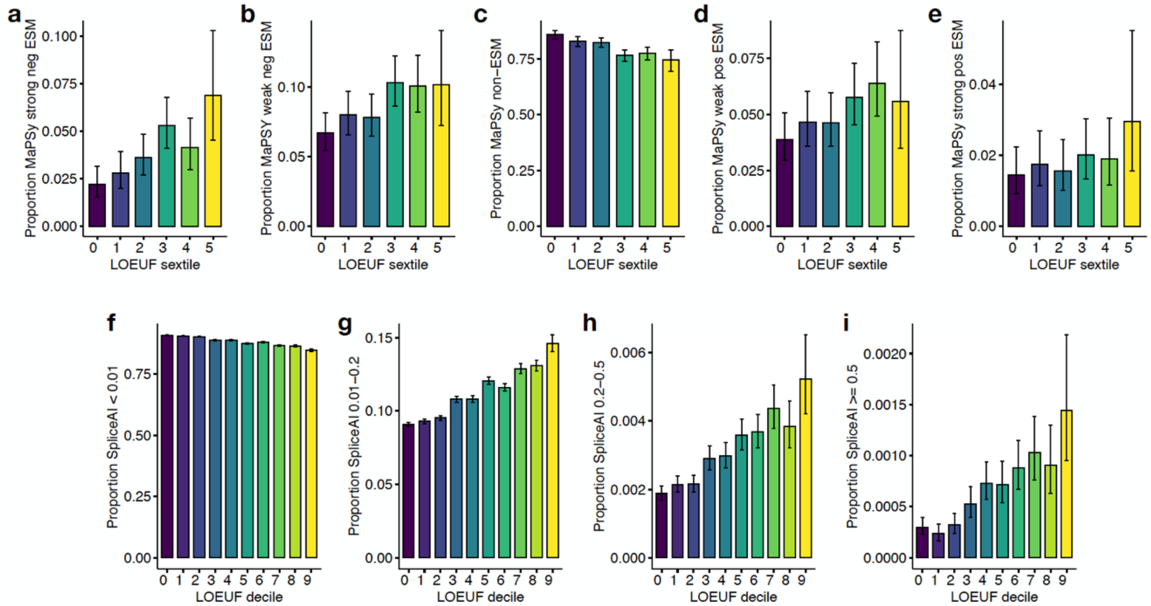


**Fig. S6. MaPSy and SpliceAI by gene constraint. (a-e)** Proportion of assayed human evolution variants in genes of varying constraint as measured by loss-of-function observed/expected upper bound fraction (LOEUF) sextiles that were identified by MaPSy as strong negative ESM (FC<-3), weak negative ESM (-3< FC<-1.5), non-ESM (-1.5<FC<1.5), weak positive ESM (1.5<FC<3) or strong positive ESM (FC>3). **(f-i)** Proportion of human evolution variants in LOEUF deciles that were identified by SpliceAI as non-splice disrupting (<0.01), weak (0.01-0.2), moderate (0.2-0.5), or strong (>0.5).

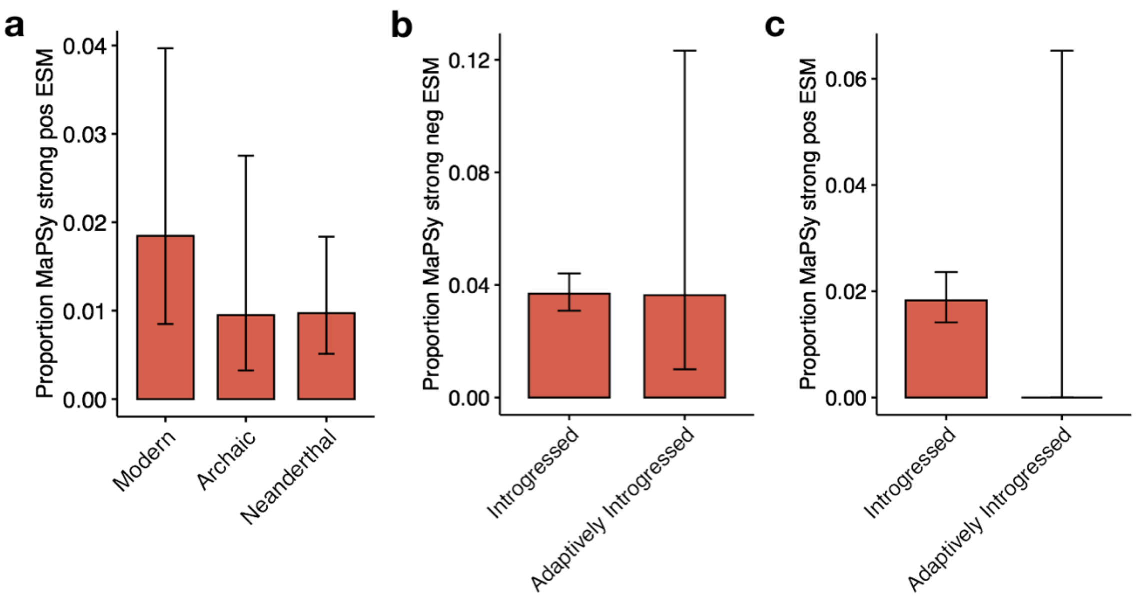


**Fig. S7. Non-significant differences in MaPSy ESM proportions between human evolution variant sets. (a)** Differences in strong positive (FC > 3) ESM proportions between modern-, archaic-, and Neanderthal-specific variants. Differences in **(b)** strong negative (FC < -3) or **(c)** strong positive (FC > 3) proportions between introgressed and adaptively introgressed variants.

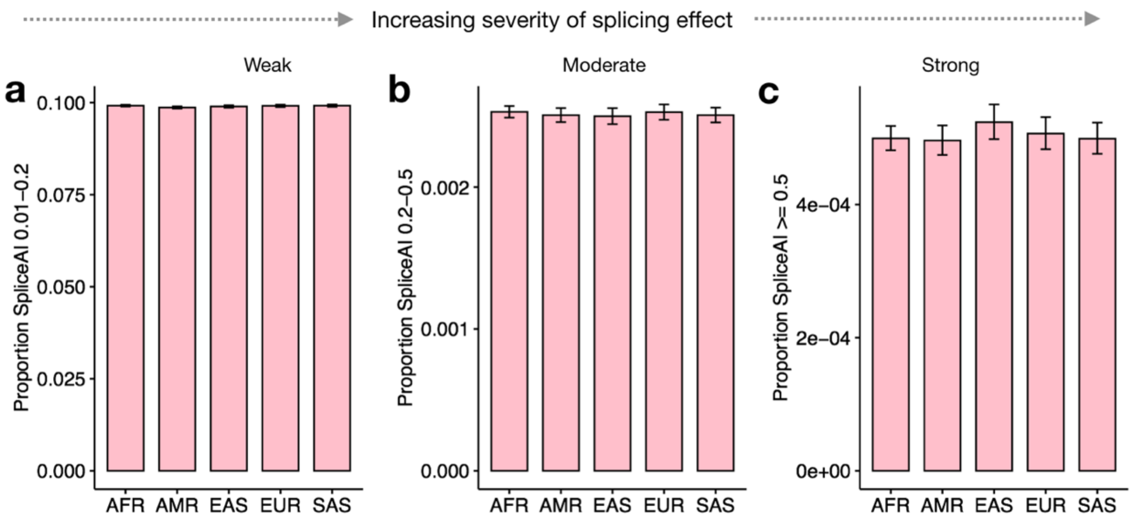


**Supplementary Table S8. No differences in SpliceAI scores for variants with MAF ≥ 0.01 between 1KGP super-populations.** Variants were binned by SpliceAI max raw scores into no splicing effect (not shown), **(a)** weak splicing effect (between 0.01 and 0.2), **(b)** moderate splicing effect (0.2-0.5), and **(c)** strong splicing effect (0.5-1). Variant sets were compared based on the proportion of common or low frequency variants (MAF ≥ 0.01) in SpliceAI bins found in 1KGP African (AFR), American (AMR), East Asian (EAS), European (EUR), and South Asian (SAS) super-populations.

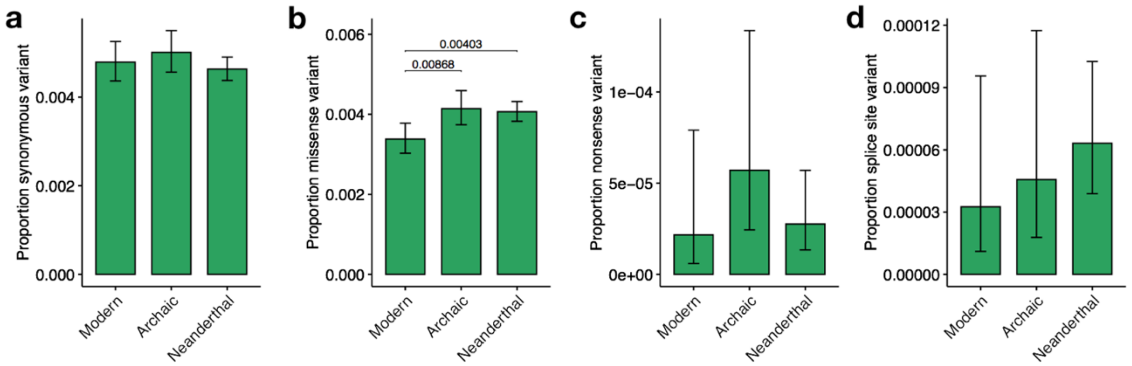


**Fig. S9. Differences in proportion of VEP consequences between lineage-specific variants.** Differences in proportion of **(a)** synonymous, **(b)** missense, **(c)** nonsense, and **(d)** splice site annotations between modern-, archaic-, and Neanderthal-specific variants. Pairwise P-values from Fisher's exact tests, p-values < 0.1 shown.

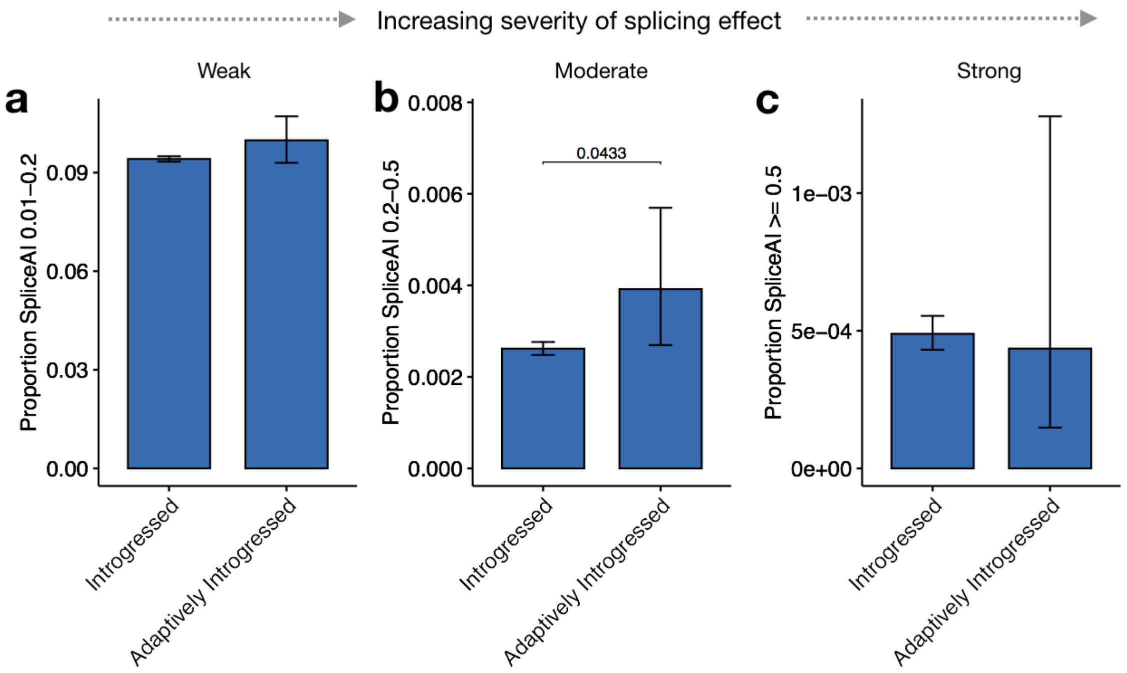


**Fig. S10. Differences in proportions of SpliceAI scores between introgressed and adaptively introgressed variants.** Variants were binned by SpliceAI max raw scores into no splicing effect (not shown), **(a)** weak splicing effect (between 0.01 and 0.2), **(b)** moderate splicing effect (0.2-0.5), and **(c)** strong splicing effect (0.5-1). Variant sets were compared based on the proportion of variants in SpliceAI bins between introgressed and adaptively introgressed variants. Pairwise P-values from Fisher's exact tests, p-values < 0.1 shown.

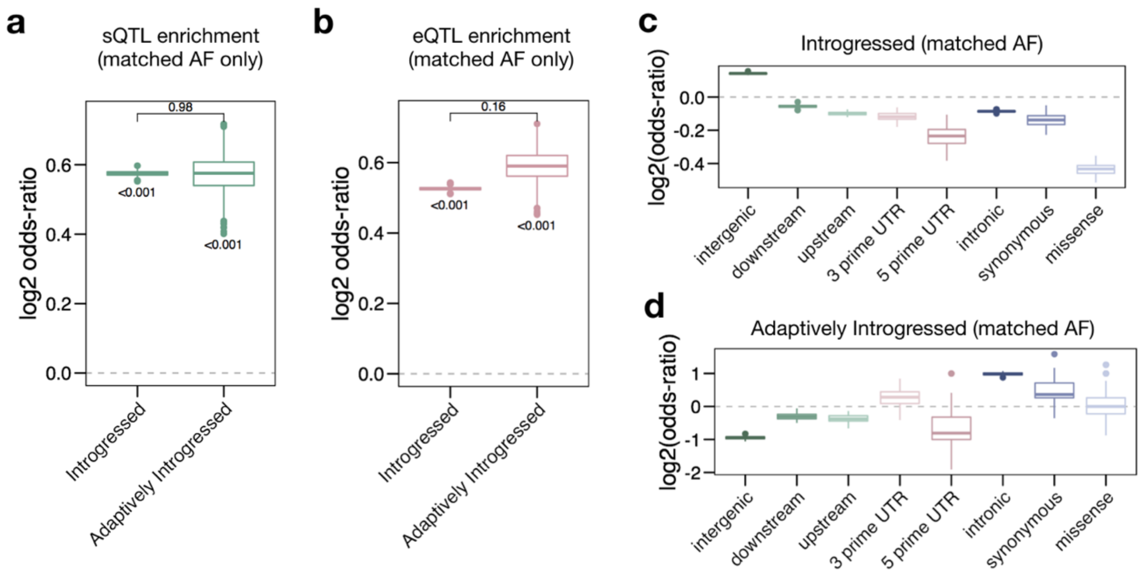


**Fig. S11. Enrichment of GTEx QTLs among introgressed and adaptively introgressed variants. (a,b)** Enrichment of GTEx sQTLs and eQTLs significant at FDR < 0.05 in at least one tissue among introgressed and adaptively introgressed variants using AF but not LD matched controls. Variability of estimates, one-sample p-values, and pairwise p-values are based on 1,000 randomly sampled sets. See Materials and Methods for details on p-value calculations. **(c,d)** Enrichment (log2 odds-ratio) of VEP consequences among introgressed and adaptively introgressed variants for AF matched controls.

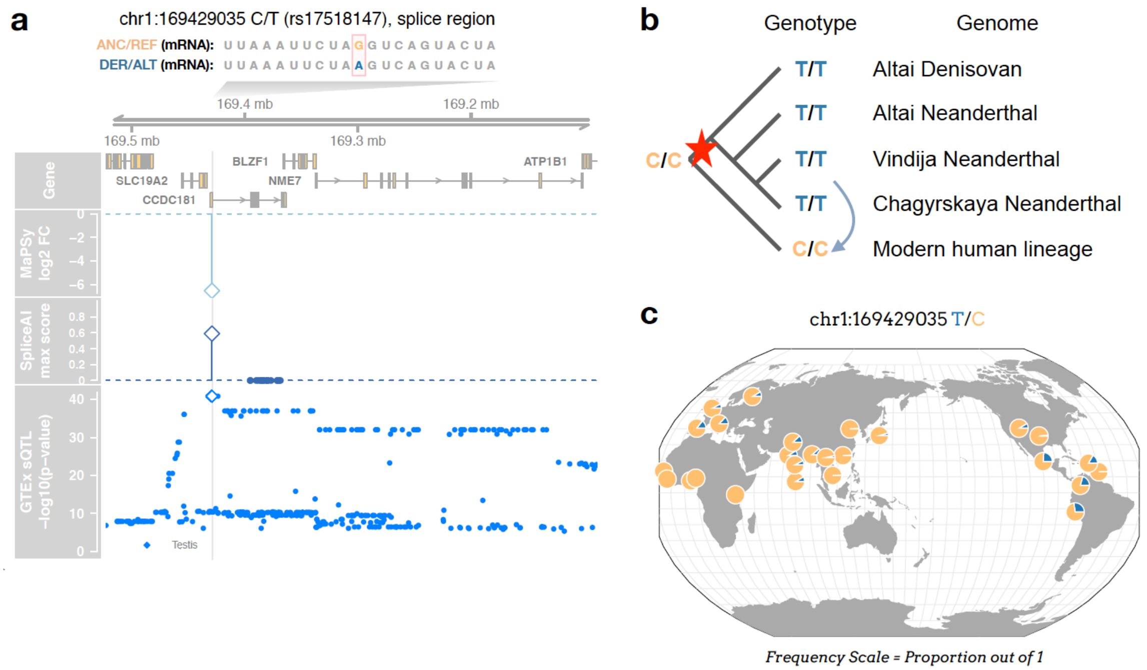


**Fig. S12. Archaic-specific and introgressed splice region ESM (rs17518147) in *CCDC181*.** **(a)** Genome tracks showing MaPSy, SpliceAI, and GTEx sQTL results for rs10128298 (shown as diamonds) and nearby introgressed variants. **(b)** Comparisons between genotypes in four high-coverage archaic genomes and the consensus modern human genotype (red star represents origin of variant, light blue arrow represents introgression event). **(c)** 1KGP allele frequencies across 26 populations from the Geography of Genetic Variants Browser.

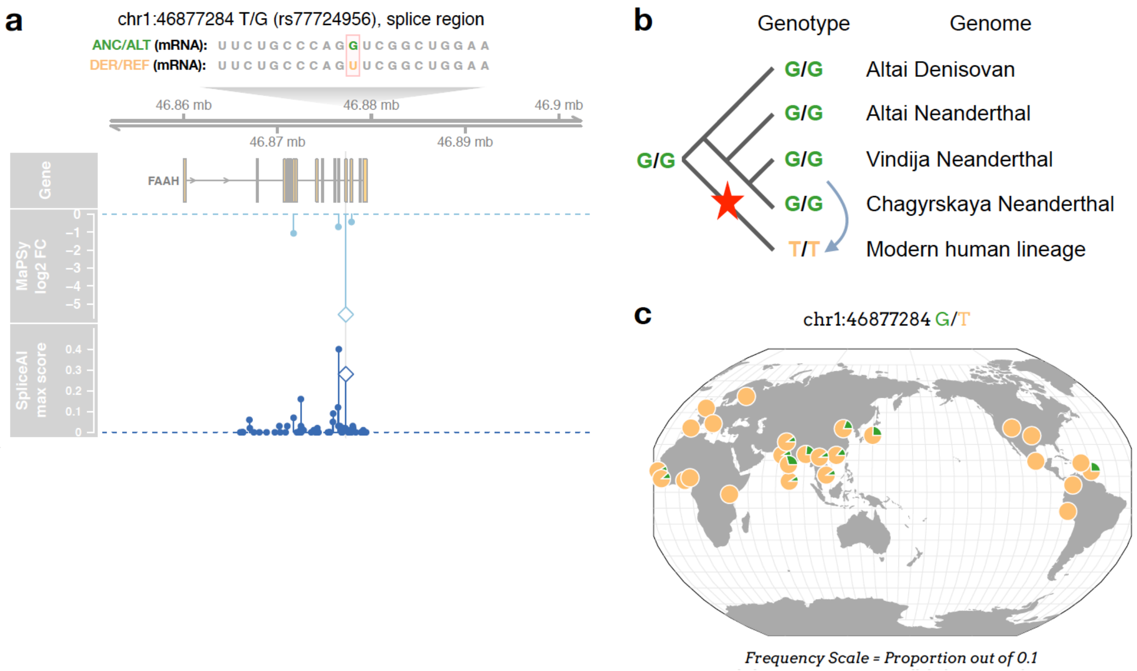


**Fig. S13. Nearly-fixed modern-specific splice region ESM with an introgressed reintroduction of an ancestral allele (rs77724956) in *FAAH*. (a)** Genome tracks showing MaPSy and SpliceAI results for rs77724956 (shown as diamonds) and nearby introgressed variants. **(b)** Comparisons between genotypes in four high-coverage archaic genomes and the consensus modern human genotype (red star represents origin of variant, light blue arrow represents an introgression event). **(c)** 1KGP allele frequencies across 26 populations from the Geography of Genetic Variants Browser.

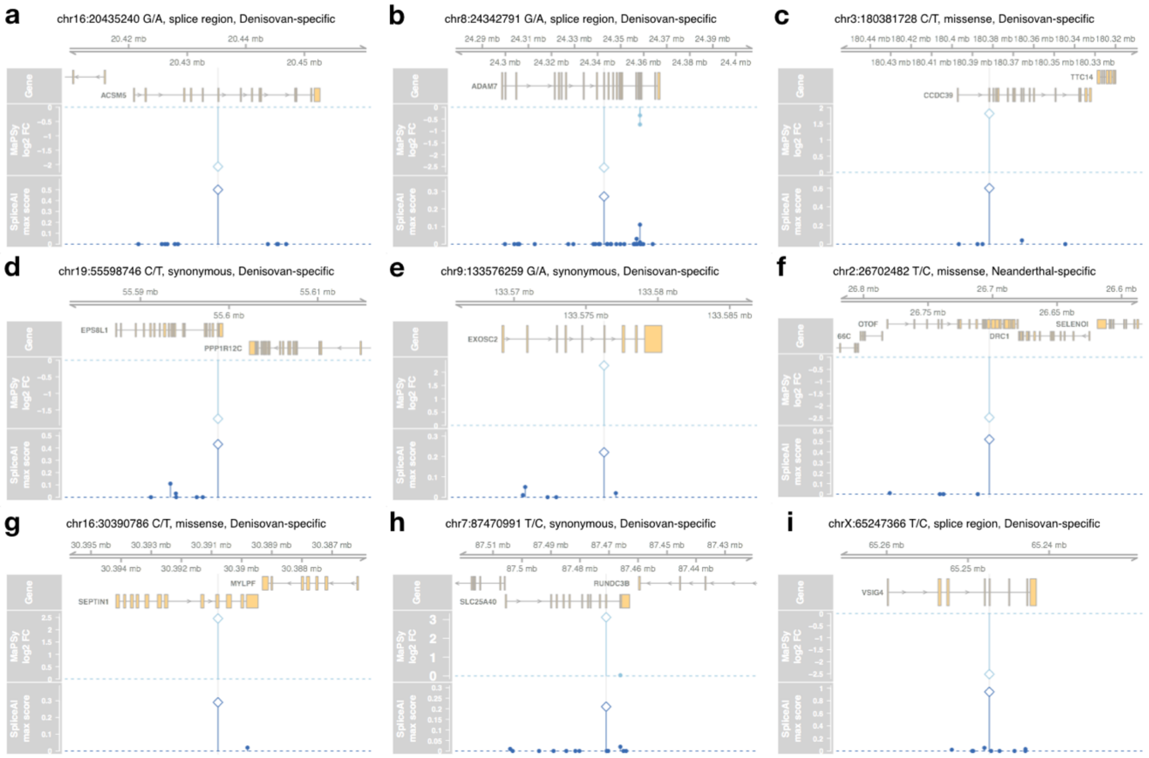


**Fig. S14. Neanderthal and Denisovan-specific variants not found in 1KGP or gnomAD. (a-i)** Genome tracks showing MaPSy and SpliceAI results for private splicing variants that are MaPSy strong ESMs with SpliceAI max raw score >= 0.2 and which are not introgressed into modern humans or polymorphic in 1KGP or gnomAD genomes in *ACSM5*, *ADAM7*, *CCDC39*, *EPS8L1*, *EXOSC2*, *OTOF*, *SEPTIN1*, *SLC25A40*, and *VSIG4*. Results for all 36 private splicing variants that are MaPSy ESMs with SpliceAI max raw scores >= 0.1 are listed in Supplementary Data S8.

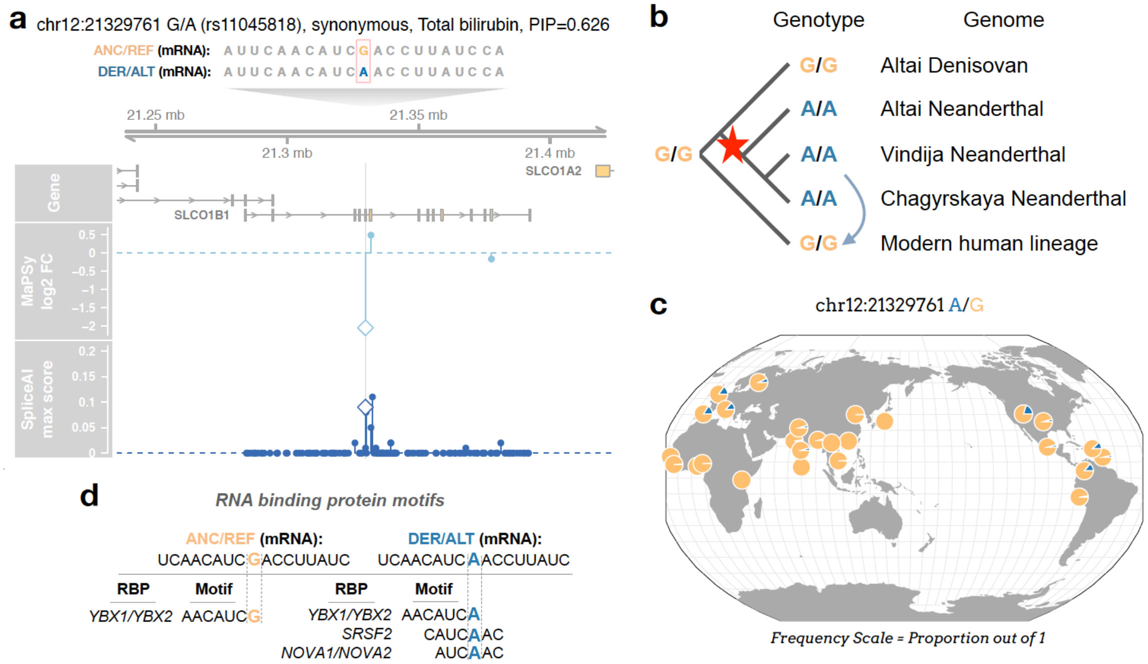


**Fig. S15. Neanderthal-derived introgressed synonymous ESM (rs11045818) overlapping a fine-mapped total bilirubin-associated variant in *SLCO1B1*. (a)** Genome tracks showing MaPSy and SpliceAI results for rs11045818 (shown as diamonds) and nearby introgressed variants. **(b)** Comparisons between genotypes in four high-coverage archaic genomes and the consensus modern human genotype (red star represents origin of variant, light blue arrow represents an introgression event). **(c)** 1KGP allele frequencies across 26 populations from the Geography of Genetic Variants Browser. **(d)** RNA binding protein (RBP) motifs (min width of 6) identified in the ANC or DER sequences using ATtRACT.

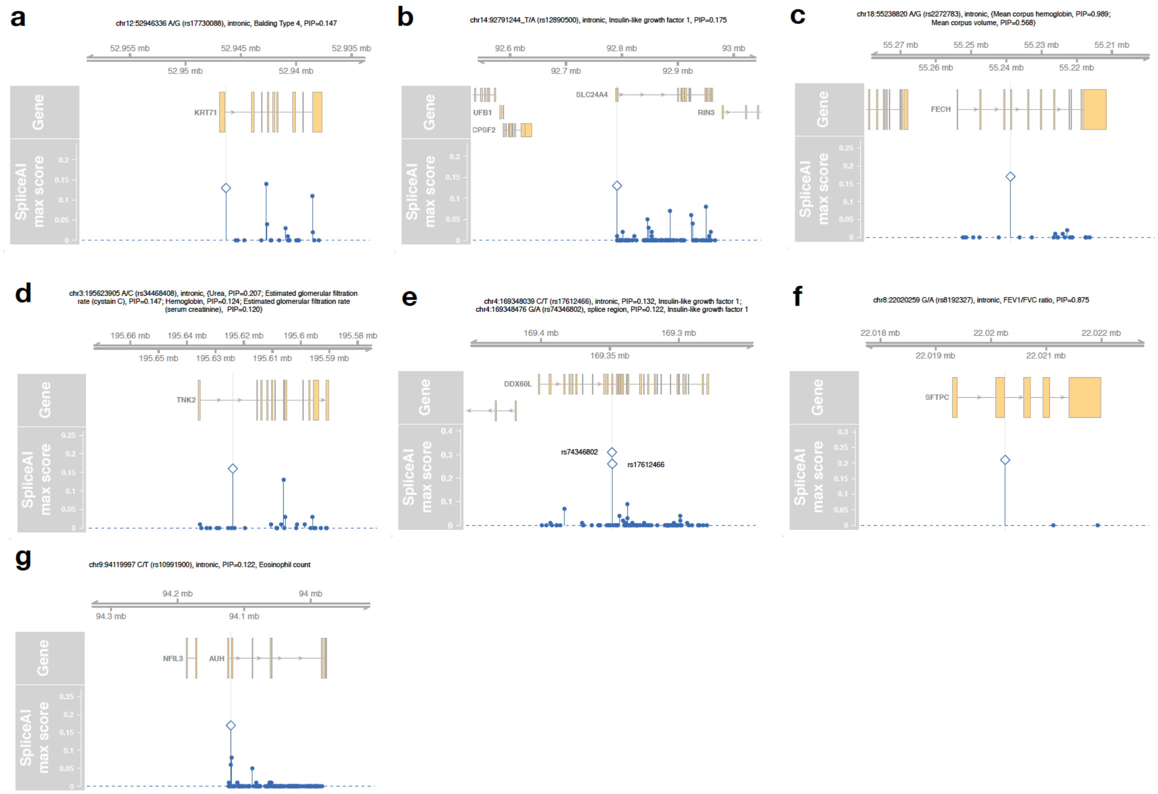

**Fig. S16. Overlapping hits between UK Biobank fine-mapped variants with PIP > 0.1 and human evolution variants with SpliceAI max > 0.1.** Genome tracks showing SpliceAI results for: **(a)** an adaptively introgressed variant in the keratin gene *KRT71*; and **(b-g)** archaically introgressed variants in *SLC24A4*, *FECH*, *TNK2*, two unique variants in *DDX60L, SFTPC*, and *AUH*.
